# The relationship between hippocampal-dependent task performance and hippocampal grey matter myelination and iron content

**DOI:** 10.1101/2020.08.18.255992

**Authors:** Ian A. Clark, Martina F. Callaghan, Nikolaus Weiskopf, Eleanor A. Maguire

**Author notes:** Corresponding author: Eleanor A. Maguire.

## Abstract

Individual differences in scene imagination, autobiographical memory recall, future thinking and spatial navigation have long been linked with hippocampal structure in healthy people, although evidence for such relationships is, in fact, mixed. Extant studies have predominantly concentrated on hippocampal volume. However, it is now possible to use quantitative neuroimaging techniques to model different properties of tissue microstructure in vivo such as myelination and iron. Previous work has linked such measures with cognitive task performance, particularly in older adults. Here we investigated whether performance on scene imagination, autobiographical memory, future thinking and spatial navigation tasks was associated with hippocampal grey matter myelination or iron content in young, healthy adult participants. MRI data were collected using a multi-parameter mapping protocol (0.8mm isotropic voxels) from a large sample of 217 people with widely-varying cognitive task scores. We found little evidence that hippocampal grey matter myelination or iron content was related to task performance. This was the case using different analysis methods (voxel-based quantification, partial correlations), when whole brain, hippocampal regions of interest, and posterior:anterior hippocampal ratios were examined, and across different participant sub-groups (divided by gender, task performance). Variations in hippocampal grey matter myelin and iron levels may not, therefore, help to explain individual differences in performance on hippocampal-dependent tasks, at least in young, healthy individuals.

## Introduction

Variations in hippocampal structure within the healthy population have long been posited to influence performance on tasks known to be hippocampal-dependent, such as scene imagination, autobiographical memory recall, future thinking and spatial navigation. Extant studies have predominantly examined this relationship in terms of hippocampal grey matter volume. However, in reviewing the literature, Clark et al. (2020) found mixed evidence for an association between hippocampal grey matter volume and performance on tasks assessing these cognitive functions in healthy individuals. They then proceeded to examine this issue in-depth by collecting data from a large sample of 217 young, healthy, adult participants, but found little evidence that hippocampal grey matter volume was related to task performance (Clark et al., 2020).

It could be argued, however, that hippocampal grey matter volume is too blunt an instrument to consistently detect structure-function relationships in healthy young people. By contrast, it is now possible to use quantitative neuroimaging techniques to model different properties of tissue microstructure, such as myelination and iron content (Weiskopf et al., 2015). Myelin in brain tissue is thought to facilitate connectivity between neurons, with greater levels of myelination increasing the speed with which neurons can communicate (Nave and Werner, 2014). Iron is also important to consider as it is involved in the production and maintenance of myelin, and is therefore required for normal brain function (Mills et al., 2010; Todorich et al., 2009).

Myelination and iron content in grey matter can be studied in vivo in humans using a multi-parameter mapping (MPM) quantitative neuroimaging protocol (Callaghan et al., 2015; Callaghan et al., 2019; Weiskopf et al., 2013). Processing of these images using the hMRI toolbox (Tabelow et al., 2019) results in four maps that are differentially (but not solely) sensitive to specific aspects of tissue microstructure. These are: magnetisation transfer saturation (MT saturation), sensitive to myelination; proton density (PD), sensitive to tissue water content; the longitudinal relaxation rate (R_1_), sensitive to myelination, iron and water content (but primarily myelination); and the effective transverse relaxation rate (R_2_*), sensitive to tissue iron content. Extant studies have reported relationships between myelination, iron and ageing across the lifespan (Callaghan et al., 2014; Draganski et al., 2011) and in young adults (Carey et al., 2018), and also correlations with verbal memory performance in older adults (Steiger et al., 2016), and meta-cognitive ability in young adults (Allen et al., 2017). However, as far as we are aware, no studies have investigated the relationship between hippocampal grey matter myelination or iron content and scene imagination, autobiographical memory recall, future thinking or navigation ability in healthy young people. Consequently, this is what we sought to examine in the current study.

We used the large dataset (n = 217) from the Clark et al. (2020) study which comprised an MPM quantitative imaging protocol (Callaghan et al., 2015; Callaghan et al., 2019; Weiskopf et al., 2013), and cognitive task performance with wide variability in scores. While aspects of these data (hippocampal grey matter volume, cognitive task performance) have been reported before (Clark et al. 2019, 2020; Clark and Maguire 2020), the tissue microstructure MRI data, measuring myelination and iron content, have not been published previously. The mixed literature relating to hippocampal grey matter volume and the dearth of studies investigating hippocampal grey matter myelination and iron content made the formulation of clear hypotheses difficult. As such, we focussed on conducting deep and wide-ranging data analyses to characterise any links between the tissue microstructure measures and task performance in the same manner as Clark et al. (2020).

## Methods

### Participants

Two hundred and seventeen people (mean age 29.0 years ± 5.60) were recruited from the general population, 109 females and 108 males. The age range was restricted to 20-41 years old to limit the possible effects of ageing. Participants had English as their first language and reported no psychological, psychiatric or neurological health conditions. People with hobbies or vocations known to be associated with the hippocampus (e.g. licensed London taxi drivers) were excluded. Participants were reimbursed £10 per hour for taking part which was paid at study completion. All participants gave written informed consent and the study was approved by the University College London Research Ethics Committee (project ID: 6743/001).

### Procedure

Participants completed the study over three visits - structural MRI scans were acquired during the first visit, and cognitive testing was conducted during visits two and three. All participants completed all parts of the study.

### Cognitive tasks and statistical analyses

All tasks are published and were performed and scored as per their published use. Full descriptions are also provided in Clark et al. (2019, 2020) and Clark and Maguire (2020). Details of the double scoring for the current study are provided in the Supplemental Material Tables S1-S4. In brief, there were four tasks: (1) **Scene imagination** was tested using the scene construction task (Hassabis et al., 2007) which measures the ability to mentally construct visual scenes. The main outcome measure is the “experiential index”, while the sub-measures of interest are content scores and a rating of the spatial coherence of scenes. (2) **Autobiographical memory recall** was tested using the autobiographical interview (AI; Levine et al., 2002), which measures the ability to recall past experiences over four time periods from early childhood to within the last year. The two main outcome measures are the number of “internal” and “external” details. Internal details are of interest here because they describe the event in question (i.e. episodic details) and are thought to be hippocampal-dependent. Sub-measures are the separate content categories that comprise the internal details outcome measure, and also AI vividness ratings. (3) The **Future thinking** task follows the same procedure as the scene construction task, but requires participants to imagine three plausible future scenes involving themselves (an event at the weekend; next Christmas; the next time they will meet a friend). (4) **Navigation** ability was assessed using the paradigm described by Woollett and Maguire (2010). A participant watches movie clips of two overlapping routes through an unfamiliar real town four times. The main outcome measure is the combined scores from the five sub-measures used to assess navigational ability which are: movie clip recognition, recognition memory for scenes, landmark proximity judgements, route knowledge where participants place scene photographs from the routes in the correct order as if travelling through the town, and the drawing of a sketch map. Data were summarised using means and standard deviations, calculated in SPSS v22.

### MRI data acquisition and preprocessing

Three Siemens Magnetom TIM Trio MRI systems with 32 channel head coils were used to collect the structural neuroimaging data. All scanners were located at the same imaging centre, running the same software.

Whole brain images at an isotropic resolution of 800μm were obtained using a MPM quantitative imaging protocol (Callaghan et al., 2015; Callaghan et al., 2019; Weiskopf et al., 2013). This consisted of the acquisition of three multi-echo gradient acquisitions with either PD, T1 or MT weighting. Each acquisition had a repetition time, TR, of 25 ms. PD weighting was achieved with an excitation flip angle of 6^0^, which was increased to 21^0^ to achieve T1 weighting. MT weighting was achieved through the application of a Gaussian RF pulse 2 kHz off resonance with 4ms duration and a nominal flip angle of 220^0^. This acquisition had an excitation flip angle of 6^0^. The field of view was 256mm head-foot, 224mm anterior-posterior (AP), and 179mm right-left (RL). The multiple gradient echoes per contrast were acquired with alternating readout gradient polarity at eight equidistant echo times ranging from 2.34 to 18.44ms in steps of 2.30ms using a readout bandwidth of 488 Hz/pixel. Only six echoes were acquired for the MT weighted volume to facilitate the off-resonance pre-saturation pulse within the TR. To accelerate the data acquisition, partially parallel imaging using the GRAPPA algorithm was employed in each phase-encoded direction (AP and RL) with forty integrated reference lines and a speed up factor of two. Calibration data were also acquired at the outset of each session to correct for inhomogeneities in the RF transmit field (Lutti et al., 2010; Lutti et al., 2012).

Data were processed using the hMRI toolbox (Tabelow et al., 2019) within SPM12 (www.fil.ion.ucl.ac.uk/spm). The default toolbox configuration settings were used, with the exception that correction for imperfect spoiling was additionally enabled (see also Callaghan et al., 2019). As alluded to earlier, this image processing resulted in four maps which differentially modelled tissue microstructure (Figure 1): a MT saturation map sensitive to myelination, a PD map sensitive to tissue water content, a R_1_ map sensitive to myelination, iron and water content (but primarily myelination), and a R_2_* map sensitive to tissue iron content.

**Figure. 1.**
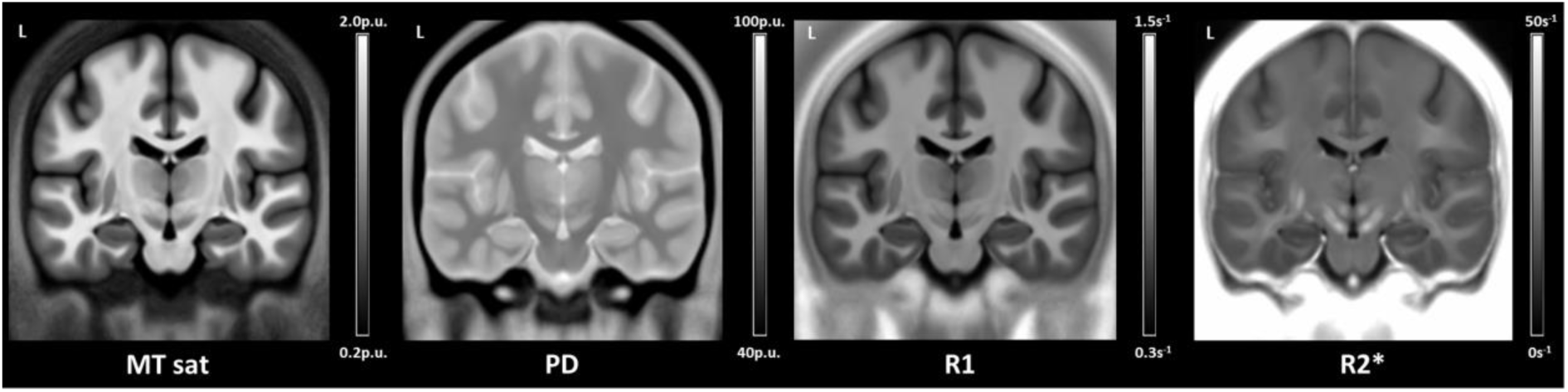
The averaged magnetisation transfer saturation (MT sat), proton density (PD), longitudinal relaxation rate (R_1_) and effective transverse relaxation rate (R_2_*) tissue microstructure maps of the whole sample (n = 217) in MNI space. The scale bars are the estimated physical values of the tissue properties in each map quantified in standardised units. For the MT saturation and PD maps this is as percent units (p.u.) and for the R_1_ and R_2_* maps this is per second (s^-1^).

Each participant’s MT saturation map was then segmented into grey and white matter probability maps using the unified segmentation approach (Ashburner and Friston, 2005), but using the tissue probability maps developed by Lorio et al. (2016) and no bias field correction (since the MT saturation map shows negligible bias field modulation). The grey and white matter probability maps were used to perform inter-subject registration using DARTEL, a nonlinear diffeomorphic algorithm (Ashburner, 2007). The resulting DARTEL template and deformations were used to normalize the MT saturation, PD, R_1_ and R_2_* maps to the stereotactic space defined by the Montreal Neurological Institute (MNI) template (at 1 × 1 × 1mm resolution), but without modulating by the Jacobian determinants of the deformation field in order to allow for the preservation of the quantitative values. Finally, a tissue weighted smoothing kernel of 4mm full width at half maximum (FWHM) was applied using the voxel-based quantification approach (VBQ; Draganski et al. 2011), which again aims to preserve the quantitative values.

### Relationships between the tissue microstructure maps

Of note, the tissue microstructure maps are not completely independent since they are estimated from the same three multi-echo gradient echo acquisitions. As such, relationships exist between the tissue microstructure maps, and a finding in one map can be used to validate a finding in another. For example, if a positive association is observed between task performance and the hippocampus in the MT saturation map, then a corresponding positive association would also be expected in the hippocampus when using the R_1_ map (since increased macromolecular content will also increase R_1_), and a corresponding negative association would be expected in the PD map (due to a reduction in free water content as the macromolecular content increases; Mezer et al., 2013).

### Primary VBQ analyses

Our analyses followed the same procedures as those detailed in Clark et al. (2020) except that here we assessed each of the tissue microstructure maps using VBQ (Draganski et al., 2011). VBQ is a similar methodology to the voxel-based morphometry (VBM) technique used to study grey matter volume (Ashburner and Friston, 2000) but one that retains the quantitative values carrying information about the tissue microstructure.

First, we examined the relationship between hippocampal grey matter in each of the tissue microstructure maps and the main outcome measure for each of the cognitive tasks assessing scene imagination, autobiographical memory, future thinking and navigation. We then examined the associations between each of the sub-measures from these tasks and hippocampal grey matter in each of the four tissue microstructure maps. Statistical analyses were carried out using multiple linear regression models with cognitive task performance as the measure of interest, while including covariates for age, gender, total intracranial volume, and the different MRI scanners. The dependent variable was the smoothed and normalised grey matter value from each tissue microstructure map.

Whole brain VBQ analyses were carried out for each tissue microstructure map using an explicitly defined mask which was generated by averaging the smoothed and normalised MT saturation grey matter probability map across all participants. When the grey matter probability was below 80%, these voxels were excluded from the analysis. Two-tailed t-tests were used, with statistical thresholds applied at p < 0.05 family-wise error (FWE) corrected for the whole brain, and a minimum cluster size of 5 voxels.

As relationships exist between the tissue microstructure maps, following the finding of a relationship in one map, we also examined whether corresponding relationships existed in the other maps, even at a more liberal statistical threshold (p < 0.001 uncorrected). Observing related associations across multiple maps was deemed supportive of a true result, while finding a correlation in only one of the tissue microstructure maps was regarded as unreliable.

### Hippocampal region of interest (ROI) VBQ

Following the whole brain analysis, we focused on the hippocampus using bilateral anatomical hippocampal masks. The masks were manually delineated on the group-averaged MT saturation map in MNI space (1 × 1 × 1 mm) using ITK-SNAP (www.itksnap.org). As in Poppenk and Moscovitch (2011) and Brunec et al. (2019), the anterior hippocampus was delineated using an anatomical mask that was defined in the coronal plane and proceeded from the first slice where the hippocampus can be observed in its most anterior extent until the final slice of the uncus. The posterior hippocampus was defined from the first slice following the uncus until the final slice of observation in its most posterior extent. The whole hippocampal mask comprised the combination of the anterior and posterior masks. Region of interest (ROI) analyses were performed using two-tailed t-tests, with voxels regarded as significant when falling below an initial whole brain uncorrected voxel threshold of p < 0.001, and then a small volume correction threshold of p < 0.05 FWE corrected for each mask, with a minimum cluster size of 5 voxels. As with the whole brain analyses, following the finding of a relationship in one map, we also examined whether corresponding relationships existed in the other maps, even at a more liberal statistical threshold of p < 0.001 uncorrected.

### Auxiliary analyses using extracted hippocampal microstructure measurements

Several auxiliary analyses were performed using the hippocampal grey matter tissue microstructure measurements that were extracted for each participant from each tissue microstructure map using ‘spm_summarise’. Whole, anterior and posterior bilateral anatomical hippocampal masks were applied to each participant’s smoothed and normalised grey matter MT saturation, PD, R_1_, and R_2_* maps, and the average value within each mask extracted. We also calculated each participant’s posterior:anterior hippocampal ratio for each tissue microstructure measurement (Poppenk and Moscovitch, 2011).

We first performed partial correlations between the extracted tissue microstructure metrics and the cognitive task performance measures. Then, we investigated the effects of gender. Next, we examined the effects of task performance by dividing the sample into low and high performing groups dependent upon the median score for each task, directly comparing the two groups. Finally we also compared the best and worst performers for each task, defined as approximately the top and bottom 10% (n ∼ 20 in each group). Age, gender, total intracranial volume and MRI scanner were included as covariates in all analyses, with the exception of those examining gender, where gender was not a covariate. As in Clark et al. (2020), statistical correction was made using false discovery rate (FDR; Benjamini and Hochberg, 1995), with a FDR of p < 0.05 allowing for 5% false positive results across the tests performed, and calculated using the resources provided by McDonald (2014).

## Results

### Cognitive task performance

A summary of the outcome measures for the cognitive tasks is shown in Table 1. A wide range of scores was obtained for all variables with the exception of navigation movie clip recognition, where performance was close to ceiling.

**Table 1.** Means and standard deviations for the main outcome measures and sub-measures of the cognitive tasks.

| Variable | Mean | Standard Deviation |
| --- | --- | --- |
| **Main outcome measures** |  |  |
| Scene construction experiential index (/60) | 40.50 | 6.08 |
| Autobiographical interview internal details (total number) | 23.95 | 7.25 |
| Future thinking experiential index (/60) | 39.12 | 7.23 |
| Navigation (/250) | 143.46 | 35.83 |
| **Sub-measures** |  |  |
| ***Scene construction*** |  |  |
| Spatial references (total number) | 3.47 | 1.56 |
| Entities present (total number) | 9.90 | 2.92 |
| Sensory descriptions (total number) | 12.25 | 3.46 |
| Thoughts/emotions/actions (total number) | 3.36 | 1.70 |
| Spatial coherence index (/6) | 2.88 | 1.57 |
| ***Autobiographical interview*** |  |  |
| Internal events (total number) | 11.02 | 3.88 |
| Internal time (total number) | 1.43 | 0.69 |
| Internal place (total number) | 2.23 | 0.63 |
| Internal perceptual (total number) | 5.91 | 3.07 |
| Internal thoughts/emotions (total number) | 3.37 | 1.69 |
| Vividness ratings (/6) | 4.62 | 0.74 |
| ***Future thinking*** |  |  |
| Spatial references (total number) | 2.53 | 1.65 |
| Entities present (total number) | 10.31 | 3.28 |
| Sensory descriptions (total number) | 8.81 | 3.59 |
| Thoughts/emotions/actions (total number) | 5.18 | 2.28 |
| Spatial coherence index (/6) | 2.63 | 1.77 |
| ***Navigation*** |  |  |
| Movie clip recognition (/16) | 15.44 | 0.86 |
| Scene recognition (/32) | 29.45 | 1.89 |
| Proximity judgements (/10) | 7.48 | 1.43 |
| Route knowledge (/24) | 11.98 | 4.62 |
| Sketch map | 79.11 | 30.91 |

### Primary VBQ analyses

As our main focus was on the relationship between cognitive task performance and hippocampal grey matter myelination and iron content, here we report findings pertaining to the hippocampus – any regions identified in the whole brain analysis that were outside the hippocampus are reported in the Supplemental Results.

No significant relationships were evident between cognitive task performance and hippocampal grey matter in any of the tissue microstructure maps. This was the case for the main outcome measures of the tasks assessing scene imagination, autobiographical memory, future thinking or navigation, and also for the sub-measures of these tasks.

### Hippocampal ROI VBQ

On examination of the hippocampal masks, no relationships were evident between cognitive task performance and hippocampal grey matter in any of the tissue microstructure maps. Considering the task sub-measures, it was either the case that they were not associated with any measure of hippocampal grey matter tissue microstructure, or the results were not validated across the tissue microstructure maps – correlations associated with only one map are reported in the Supplemental Results.

### Auxiliary analyses using extracted hippocampal microstructure measurements

The means and standard deviations of the extracted hippocampal grey matter tissue microstructure metrics are reported in the Supplemental Results, Table S5.

No relationships were identified between any of the extracted hippocampal grey matter tissue microstructure metrics and performance on any of the main or sub-measures of the cognitive tasks (see Supplemental Results Tables S6-S9 for details). These partial correlation findings therefore support those of the primary VBQ analyses.

Similarly, there were no significant effects of gender (Supplemental Results Tables S10-S17), and no significant findings from the median split direct comparisons (Supplemental Results Tables 18-22), partial correlations (Supplemental Results Tables S23-S30), and when the best and worst performers were compared (Supplemental Results Tables 31-35).

## Discussion

In this study we moved beyond hippocampal grey matter volume to examine hippocampal grey matter tissue microstructure in the form of quantitative neuroimaging biomarkers of myelination and iron content, and whether they were linked with performance on tasks known to be hippocampal-dependent. We found little evidence for any associations between these measures and scene imagination, autobiographical memory recall, future thinking and spatial navigation. This was despite having a large sample with wide-ranging scores on the cognitive tasks, using different analysis methods (voxel-based quantification, partial correlations), examining whole brain and hippocampal regions of interest (bilateral whole hippocampus, anterior and posterior portions, hippocampal posterior:anterior ratio), and different participant sub-groups (divided by gender and task performance). Variations in hippocampal grey matter myelination or iron content, seem not, therefore, to be significantly related to hippocampal-dependent task performance in young, healthy individuals.

Myelination and iron are essential for communication between neurons (Nave and Werner, 2014; Todorich et al., 2009). Therefore, it could have been that higher levels of myelin within the hippocampus would result in faster hippocampal neuron communication, enabling better task performance. Conversely, it might have been the case that higher levels of hippocampal iron were associated with lower task performance - while moderate iron levels are required for normal functioning, excessive iron accumulation can impede function (Zecca et al., 2004). Indeed, such a finding has previously been reported in young healthy adults in relation to meta-cognitive ability (Allen et al., 2017). Here, however, we found no relationships between hippocampal grey matter myelination or iron content and performance on scene imagination, autobiographical memory recall, future thinking or spatial navigation tasks, aligning with previous null results relating to hippocampal grey matter volume more generally (Clark et al., 2020; Maguire et al., 2003; Van Petten, 2004; Weisberg et al., 2019).

Null results can, of course, be difficult to interpret, and an absence of evidence is not necessarily evidence of absence. However, we believe the depth and breadth of our analyses permit confidence in these results. First, we had a large sample size with wide variance on our measures of interest, both in terms of task performance (with the exception of the navigation movie clip recognition sub-measure where performance was close to ceiling), and the tissue microstructure metrics. We were, therefore, well-placed to identify potential relationships between hippocampal grey matter myelination and iron content and cognitive task performance. Second, we used two different analysis methodologies, as well as employing bilateral hippocampal ROIs, and even divided the sample into multiple subgroups. Regardless of these different approaches to data analysis, no reliable associations between hippocampal grey matter myelination or iron content and task performance were identified. The results, therefore, do not seem to be consequent upon a specific analysis technique or group of participants, but instead there was a consistent pattern of null results.

That is not to say that relationships between hippocampal grey matter myelination or iron content and performance on our cognitive tasks of interest may not exist in other contexts. In older participants, for example, associations between hippocampal grey matter volume and performance on autobiographical and navigation tasks have been identified (e.g. Hedden et al., 2014; Moffat et al., 2006), despite less reliable evidence for such relationships in healthy young adults (Clark et al., 2020; Maguire et al., 2003; Van Petten, 2004; Weisberg et al., 2019). Changes in the extent of grey matter myelination and iron content have been documented with ageing (e.g. Callaghan et al., 2014; Carey et al., 2018; Draganski et al., 2011), as has a link between poorer verbal memory performance, decreased myelin content and increased levels of iron in the ventral striatum in older adults (Steiger et al., 2016). Therefore, similar investigations to those performed here using an older sample could reveal associations not observed in the current study.

Similarly, relationships between hippocampal grey matter myelination or iron content and task performance might only be apparent in special populations. The extreme spatial navigation of licensed London taxi drivers who have acquired “The Knowledge” of London’s layout has been reliably associated with increased posterior, but decreased anterior, hippocampal grey matter volume (e.g. Maguire et al., 2000; Maguire et al., 2006; Woollett et al., 2008; Woollett and Maguire, 2011). Variations may also exist, therefore, in hippocampal grey matter myelination and iron content, and these differences may help to elucidate how a larger posterior hippocampus might be beneficial to London taxi drivers. Comparisons between healthy individuals and patients with memory loss may also reveal important differences in the extent of myelination and iron content in the hippocampus but also elsewhere in the brain.

The hippocampus is composed of distinct subfields, and myelination or iron content in specific subregions may be associated with task performance, as opposed to associations being apparent at the level of the whole hippocampus. Some evidence exists to suggest this may be the case for hippocampal grey matter volume (Chadwick et al., 2014; Palombo et al., 2018), and future studies could investigate relationships between task performance and subfield-specific hippocampal grey matter tissue microstructure.

It is also possible to obtain different measures of tissue microstructure to those studied here. Using diffusion MRI, the fractional anisotropy (FA) of right hippocampal grey matter has been associated with navigation ability (Iaria et al., 2008), and the FA and mean diffusivity of various white matter pathways, including the fornix, have been linked with autobiographical memory, future thinking and navigation performance (Hodgetts et al., 2017; Hodgetts et al., 2020; Irish et al., 2014; Metzler-Baddeley et al., 2011; Williams et al., 2020). While replication of these results in larger samples is required, examining these tissue microstructure metrics, or white matter instead of grey matter, may reveal relationships between brain tissue microstructure and performance that were not evident in the current study. Moreover, by combining diffusion MRI and the quantitative neuroimaging techniques used here additional, more biologically interpretable microstructure measurements, such as the axonal g-ratio (e.g. Mohammadi et al., 2015), can be obtained, opening up further avenues of investigation to understand the neural basis of individual differences in cognition.

In conclusion, having tested a large sample of young healthy participants with a wide range of scores on multiple cognitive tasks, no credible associations between hippocampal grey matter myelination or iron content and scene imagination, autobiographical memory, future thinking and spatial navigation performance were identified. Consequently, variability in hippocampal grey matter myelination or iron content seem unlikely to explain individual differences in ability, at least in the general population of young healthy people. Further investigation is required to establish whether such measures may be informative in other cohorts, as well as combining myelination and iron content with other tissue microstructure techniques to examine the basis of individual differences in key aspects of cognition including recalling the past and imagining the future.

## Acknowledgements

Thanks to Anna Monk, Victoria Hotchin, Gloria Pizzamiglio and Alice Liefgreen for assistance with data collection and scoring.

## Author contributions

**Ian A. Clark:** Conceptualization, Methodology, Investigation, Formal analysis, Writing - original draft. **Martina F. Callaghan:** Formal analysis, Writing - review & editing. **Nikolaus Weiskopf:** Resources, Writing - review & editing. **Eleanor A. Maguire:** Conceptualization, Methodology, Formal analysis, Funding acquisition, Supervision, Writing - original draft.

## Supplemental Material

### Supplemental Methods

Double scoring was performed on 20% of the cognitive data. We took the most stringent approach to identifying across-experimenter agreement. Inter-class correlation coefficients with a two-way random effects model looked for absolute agreement among the experimenters. For reference, a score of 0.8 or above is considered excellent agreement beyond chance.

**Table S1.** Double scoring of the scene construction task.

| Rating |  |  |  |  |  |
| --- | --- | --- | --- | --- | --- |
|  | Spatial<br>References | Entities<br>Present | Sensory<br>Descriptions | Thoughts/<br>Emotions/Actions | Quality<br>Ratings |
| **For each individual scene** |  |  |  |  |  |
| n = 308 | 0.90 | 0.96 | 0.94 | 0.90 | 0.90 |
| **For each individual participant (i.e. score is averaged across the seven scenes)** |  |  |  |  |  |
| n = 44 | 0.91 | 0.99 | 0.97 | 0.91 | 0.93 |

Inter-class correlation coefficients from a two way random effect model looking for absolute agreement for each content score and for the quality ratings. Four experimenters scored the whole data set (n = 217 participants, 1519 individual scenes) with double scoring performed on 20% of the data (n = 44 participants, 308 scenes) proportionally for each original experimenter.

**Table S2.** Double scoring of the autobiographical interview.

| Rating |  |  |  |  |  |  |
| --- | --- | --- | --- | --- | --- | --- |
|  | Internal<br>Event | Internal<br>Place | Internal<br>Time | Internal<br>Perceptual | Internal<br>Emotion | Internal<br>Sum |
| **For each individual memory** |  |  |  |  |  |  |
| n = 215 | 0.92 | 0.85 | 0.94 | 0.92 | 0.86 | 0.94 |
| **For each individual participant (i.e. score is averaged across the five memories)** |  |  |  |  |  |  |
| n = 43 | 0.95 | 0.88 | 0.96 | 0.94 | 0.81 | 0.97 |

Inter-class correlation coefficients from a two way random effects model looking for absolute agreement for each score on the autobiographical interview. Three experimenters scored the whole data set (n = 217 participants, 1085 individual memories) and double scoring was performed 20% of the data (n = 43 participants, 215 individual memories) proportionally for each original experimenter.

**Table S3.** Double scoring of the future thinking task.

| Rating |  |  |  |  |  |
| --- | --- | --- | --- | --- | --- |
|  | Spatial<br>References | Entities<br>Present | Sensory<br>Descriptions | Thoughts/<br>Emotions/Actions | Quality<br>Ratings |
| **For each individual scene** |  |  |  |  |  |
| n = 132 | 0.90 | 0.94 | 0.93 | 0.88 | 0.90 |
| **For each individual participant (i.e. score is averaged across the three future scenes)** |  |  |  |  |  |
| n = 44 | 0.94 | 0.95 | 0.96 | 0.88 | 0.92 |

Inter-class correlation coefficients from a two way random effects model looking for absolute agreement for each content score and for the quality ratings. Four experimenters scored the whole data set (n = 217 participants, 651 individual future scenes) with double scoring performed on 20% of the data (n = 44 participants, 132 future scenes) proportionally for each original experimenter.

**Table S4.**
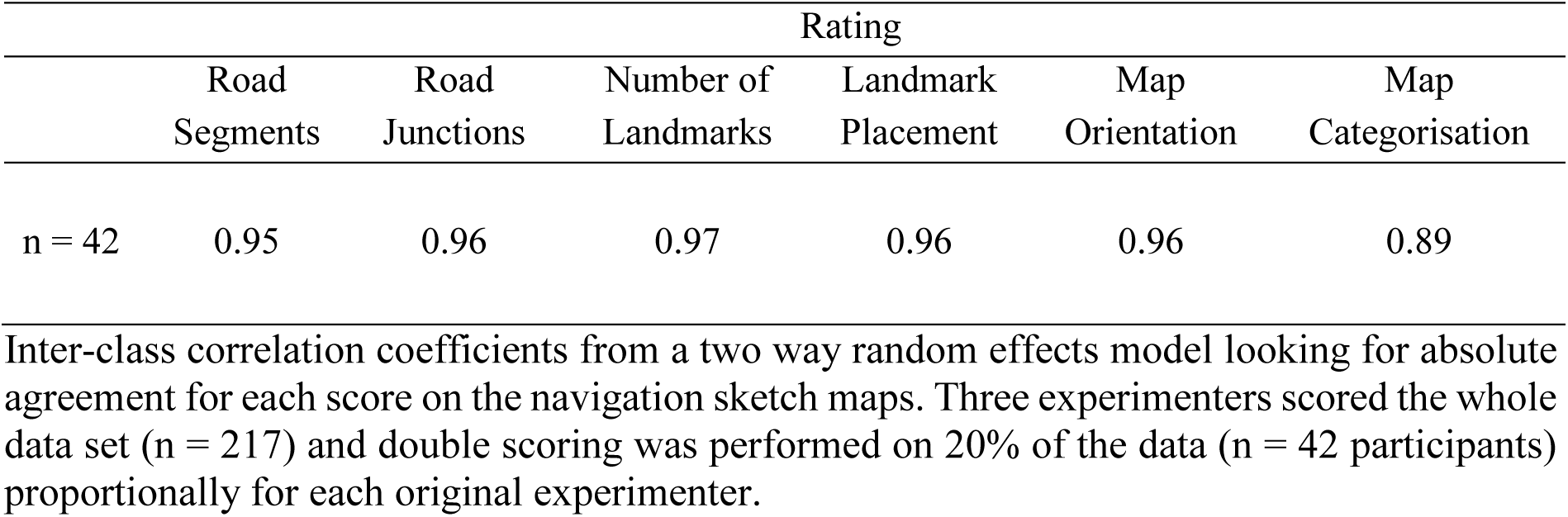
Double scoring of the navigation sketch maps.

### Supplemental Results

#### Primary analyses: VBQ results outside of the hippocampus

A small number of relationships were identified between cognitive task performance and the grey matter tissue microstructure maps outside of the hippocampus (when using a statistical threshold of p < 0.05 FWE whole brain corrected).

Within the scene construction outcome measures, one potential relationship was observed; a positive association between scene construction sensory details and 15 voxels in the left middle occipital cortex in the PD map (peak coordinates = −42 −87 17, peak t = 5.30, p_FWE whole brain corrected_ = 0.018). However, no corresponding negative associations with MT or R_1_, nor any corresponding relationships with R_2_*, were identified, even when reducing the statistical threshold to p < 0.001 uncorrected.

For future thinking, two potential associations were found. First, a positive relationship was observed in the PD map between 36 voxels in the right superior occipital gyrus and the future thinking experiential index (peak coordinates = 25 −90 33, peak t = 5.51, p_FWE whole brain corrected_ = 0.007). Reducing the statistical threshold to p < 0.001 uncorrected identified a corresponding negative association between the right superior occipital gyrus and the future thinking experiential index in the R_2_* map (cluster size = 808, peak coordinates = 25 −89 35, peak t = 4.53, p_uncorrected_ < 0.001). Overall, higher water content in the right superior occipital gyrus seems to be associated with greater future thinking experiential index scores.

Second, a positive correlation was found in the R_1_ map between 5 voxels in the left middle cingulate cortex and the future thinking experiential index (peak coordinates = 0 −20 45, peak t = 5.19, p_FWE whole brain corrected_ = 0.031). Reducing the statistical threshold to p < 0.001 uncorrected identified a corresponding negative association between the left middle cingulate cortex and the future thinking experiential index in the PD map (cluster size = 28, peak coordinates = −1, −20, 45 peak t = 3.77, p_uncorrected_ < 0.001). An increase in macromolecular content and corresponding reduction in free water content in the middle cingulate cortex, may, therefore, be associated with greater future thinking experiential index scores.

Considering navigation, for the navigation movie clip recognition task, two clusters were identified on the edge of the right occipital pole in both the PD and the R_2_* maps. In the PD map, these relationships were positive (Cluster 1: size = 7 voxels, peak coordinates = 32 − 95 15, peak t = 5.23, p = 0.024; Cluster 2: size = 10 voxels, peak coordinates = 26 −97 20, peak t = 5.21, p = 0.026), while in the R_2_* map the corresponding negative relationships were observed (Cluster 1: size = 40 voxels, peak coordinates = 32 −95 15, peak t = 5.55, p = 0.006; Cluster 2: size = 18 voxels, peak coordinates = 25 −98 19, peak t = 5.48, p = 0.009). A decrease in iron and corresponding increase in free water content in the right occipital pole, may, therefore, be related to higher navigation movie clip recognition performance.

For the navigation scene recognition task, there was a positive relationship between performance and 36 voxels in the right cuneus in the R_2_* map (peak coordinates = 17 −60 10, peak t = 5.83, p _FWE whole brain corrected_ = 0.002). However, no corresponding associations in the MT saturation or PD or R_1_ maps, were identified in the right cuneus, even when reducing the statistical threshold to p < 0.001 uncorrected.

#### Hippocampal ROI VBQ: significant results that were not validated across the tissue microstructure maps

Within the scene construction sub-measures one potential relationship was identified. A negative association was found between the scene construction spatial coherence index and 77 voxels in the left hippocampus in the PD map when using the bilateral posterior hippocampal mask (peak coordinates = −19 −41 4, peak z = 3.79, p _FWE posterior hippocampus ROI corrected_ = 0.033). However, this relationship was not significant when correcting for the bilateral whole hippocampal mask (p _FWE whole hippocampus ROI corrected_ = 0.054). In addition, no corresponding associations were identified between the scene construction spatial coherence index scores and the hippocampus in the MT saturation, R_1_ or R_2_* maps, even when reducing the statistical threshold to p < 0.001 uncorrected.

Within the autobiographical memory sub-measures, three potential associations were observed. First, a positive association was identified between AI emotion and a cluster of 45 voxels in the left hippocampus in the PD map when using the bilateral anterior hippocampal mask (peak coordinates = −26 −9 −27, peak z = 3.64, p _FWE anterior hippocampus ROI corrected_ = 0.033). However, this relationship was not significant when correcting for the whole hippocampus mask (p _FWE whole hippocampus ROI corrected_ = 0.073). In addition, no corresponding associations were observed in the hippocampus when using the MT saturation R_1_ or R_2_*maps, even when reducing the statistical threshold to p < 0.001 uncorrected.

Second, a negative association was observed between AI vividness ratings and a cluster of 37 voxels in the right hippocampus in the MT saturation map when using the bilateral whole hippocampal mask (peak coordinates = 34 −20 −12, peak z = 4.11, p _FWE whole hippocampus ROI corrected_ = 0.018), split approximately equally between the anterior and posterior hippocampal masks (anterior cluster: cluster size = 21 voxels, peak coordinates = 34 −20 −12, peak z = 4.11, p _FWE anterior hippocampus ROI corrected_ = 0.008; posterior cluster: cluster size = 16 voxels, peak coordinates = 34 −21 −12, peak z = 3.87, p _FWE posterior hippocampus ROI corrected_ = 0.027). However, no corresponding associations with the hippocampus were found in the R_1_, PD or R_2_* maps, even when reducing the statistical threshold to p < 0.001 uncorrected.

AI vividness was also negatively associated with a cluster of 56 voxels in the left hippocampus in the PD map following correction for the bilateral whole hippocampal mask (peak coordinates = −28 −27 −9, peak z = 4.12, p _FWE whole hippocampus ROI corrected_ = 0.014), localised to the posterior hippocampus (p _FWE posterior hippocampus ROI corrected_ = 0.008). However, no corresponding associations were found between AI vividness and the hippocampus in the MT saturation, R_1_ or R_2_* maps, even when reducing the statistical threshold to p < 0.001 uncorrected.

Within the future thinking sub-measures, two potential relationships were identified. A positive association was observed between future thinking spatial references and 33 voxels in the left hippocampus in the R_1_ map when using the bilateral posterior hippocampal mask (peak coordinates = −18 −22 −21, peak z = 3.75, p _FWE posterior hippocampus ROI corrected_ = 0.032). However, this relationship was not significant when correcting for the bilateral whole hippocampus mask (p _FWE whole hippocampus ROI corrected_ = 0.052). Furthermore, no corresponding associations were observed in the hippocampus in any of the other tissue microstructure maps, even when reducing the statistical threshold to p < 0.001 uncorrected.

Second, a negative association was found between the future thinking spatial coherence index and a cluster of 54 voxels in the right posterior hippocampus in the MT saturation map when using the bilateral posterior hippocampus mask (peak coordinates = 35, −29, −13, peak z = 3.76, p _FWE posterior hippocampus ROI corrected_ = 0.039). However, this relationship was not significant when correcting for the bilateral whole hippocampus mask (p _FWE whole hippocampus ROI corrected_ = 0.064) and the corresponding associations were not observed in the hippocampus when using any of the other tissue microstructure maps, even when reducing the statistical threshold to p < 0.001 uncorrected.

**Table S5.** Means and standard deviations of the extracted microstructure measurements for the hippocampal ROIs.

| Tissue<br>microstructure<br>map | Whole<br>hippocampus |  | Anterior<br>hippocampus |  | Posterior<br>hippocampus |  | Posterior/Anterior<br>ratio |  |
| --- | --- | --- | --- | --- | --- | --- | --- | --- |
|  | Mean | SD | Mean | SD | Mean | SD | Mean | SD |
| MT | 0.88 | 0.049 | 0.85 | 0.057 | 0.90 | 0.049 | 1.07 | 0.045 |
| PD | 79.87 | 1.08 | 79.60 | 1.45 | 80.09 | 1.01 | 1.01 | 0.015 |
| R<sub>1</sub> | 0.65 | 0.034 | 0.63 | 0.037 | 0.66 | 0.036 | 1.04 | 0.035 |
| R<sub>2</sub>* | 15.42 | 0.035 | 14.76 | 1.51 | 15.97 | 1.24 | 1.09 | 0.082 |

SD: standard deviation; MT: magnetisation transfer saturation; PD: proton density; R<sub>1</sub>: longitudinal relaxation rate; R<sub>2</sub>\*: effective transverse relaxation rate. A posterior/anterior ratio above 1 indicates greater values in the posterior relative to the anterior hippocampus.

**Table S6.** Partial correlations between task performance and hippocampal grey matter <u>MT saturation</u> with age, gender, total intracranial volume and MRI scanner as covariates.

| Performance variable | Whole hippocampus |  | Anterior hippocampus |  | Posterior hippocampus |  | Posterior/Anterior hippocampus ratio |  |
| --- | --- | --- | --- | --- | --- | --- | --- | --- |
|  | r | p | r | p | r | p | r | p |
| **Scene construction** |  |  |  |  |  |  |  |  |
| Experiential index | -0.096 | 0.59 | -0.097 | 0.59 | -0.081 | 0.59 | 0.043 | 0.74 |
| Spatial references | -0.087 | 0.59 | -0.09 | 0.59 | -0.072 | 0.63 | 0.049 | 0.73 |
| Entities present | -0.067 | 0.66 | -0.081 | 0.59 | -0.045 | 0.73 | 0.072 | 0.63 |
| Sensory descriptions | -0.12 | 0.59 | -0.14 | 0.55 | -0.084 | 0.59 | 0.11 | 0.59 |
| Thoughts/emotions/actions | -0.05 | 0.73 | -0.081 | 0.59 | -0.014 | 0.91 | 0.10 | 0.59 |
| Spatial coherence index | 0.07 | 0.63 | -0.027 | 0.83 | -0.10 | 0.59 | -0.085 | 0.59 |
| **Autobiographical interview** |  |  |  |  |  |  |  |  |
| Internal details | -0.005 | 0.99 | 0.002 | 0.99 | -0.012 | 0.99 | -0.026 | 0.99 |
| Internal events | 0.009 | 0.99 | 0.012 | 0.99 | 0.005 | 0.99 | -0.016 | 0.99 |
| Internal time | 0.05 | 0.99 | 0.095 | 0.85 | 0.001 | 1.00 | -0.15 | 0.75 |
| Internal place | -0.066 | 0.93 | -0.04 | 0.99 | -0.083 | 0.88 | -0.057 | 0.99 |
| Internal perceptual | -0.006 | 0.99 | -0.015 | 0.99 | 0.003 | 0.99 | 0.02 | 0.99 |
| Internal thoughts/emotions | -0.028 | 0.99 | -0.013 | 0.99 | -0.039 | 0.99 | -0.032 | 0.99 |
| Vividness rating | -0.10 | 0.85 | -0.092 | 0.85 | -0.10 | 0.85 | 0.007 | 0.99 |
| **Future thinking** |  |  |  |  |  |  |  |  |
| Experiential index | -0.14 | 0.31 | -0.13 | 0.31 | -0.13 | 0.31 | 0.05 | 0.81 |
| Spatial references | -0.15 | 0.31 | -0.14 | 0.31 | -0.14 | 0.31 | 0.05 | 0.81 |
| Entities present | -0.038 | 0.87 | -0.032 | 0.90 | -0.038 | 0.87 | 0.015 | 0.97 |
| Sensory descriptions | -0.089 | 0.51 | -0.11 | 0.38 | -0.061 | 0.70 | 0.10 | 0.38 |
| Thoughts/emotions/actions | -0.021 | 0.94 | -0.043 | 0.87 | 0.002 | 0.99 | 0.049 | 0.81 |
| Spatial coherence index | -0.10 | 0.38 | -0.062 | 0.70 | -0.13 | 0.31 | -0.062 | 0.70 |
| **Navigation** |  |  |  |  |  |  |  |  |
| Overall navigation score | -0.11 | 0.91 | -0.071 | 0.96 | -0.14 | 0.91 | -0.058 | 0.96 |
| Movie clip recognition | 0.041 | 0.99 | 0.06 | 0.96 | 0.018 | 0.99 | -0.057 | 0.96 |
| Scene recognition | 0.07 | 0.96 | 0.071 | 0.96 | 0.059 | 0.96 | -0.033 | 0.99 |
| Proximity judgements | -0.076 | 0.96 | -0.054 | 0.96 | -0.087 | 0.92 | -0.016 | 0.99 |
| Route knowledge | -0.002 | 0.99 | 0.008 | 0.99 | -0.01 | 0.99 | -0.033 | 0.99 |
| Sketch map | -0.13 | 0.91 | -0.087 | 0.92 | -0.16 | 0.91 | -0.058 | 0.96 |

P values are Benjamini-Hochberg false discovery rate corrected at $p < 0.05$ .

**Table S7.** Partial correlations between task performance and hippocampal grey matter <u>PD</u> with age, gender, total intracranial volume and MRI scanner as covariates.

| Performance variable | Whole hippocampus |  | Anterior hippocampus |  | Posterior hippocampus |  | Posterior/Anterior hippocampus ratio |  |
| --- | --- | --- | --- | --- | --- | --- | --- | --- |
|  | r | p | r | p | r | p | r | p |
| **Scene construction** |  |  |  |  |  |  |  |  |
| Experiential index | -0.028 | 0.83 | -0.044 | 0.73 | -0.001 | 0.99 | 0.053 | 0.72 |
| Spatial references | -0.025 | 0.83 | -0.078 | 0.59 | 0.046 | 0.73 | 0.13 | 0.55 |
| Entities present | -0.083 | 0.59 | -0.085 | 0.59 | -0.06 | 0.67 | 0.054 | 0.71 |
| Sensory descriptions | 0.02 | 0.86 | 0.002 | 0.99 | 0.038 | 0.77 | 0.029 | 0.82 |
| Thoughts/emotions/actions | -0.036 | 0.77 | -0.036 | 0.77 | -0.027 | 0.83 | 0.022 | 0.85 |
| Spatial coherence index | -0.042 | 0.74 | -0.004 | 0.98 | -0.078 | 0.59 | -0.06 | 0.67 |
| **Autobiographical interview** |  |  |  |  |  |  |  |  |
| Internal details | 0.045 | 0.99 | 0.093 | 0.85 | -0.026 | 0.99 | -0.14 | 0.75 |
| Internal events | 0.051 | 0.99 | 0.097 | 0.85 | -0.019 | 0.99 | -0.14 | 0.75 |
| Internal time | -0.084 | 0.88 | -0.081 | 0.88 | -0.067 | 0.93 | 0.039 | 0.99 |
| Internal place | -0.005 | 0.99 | 0.00 | 1.00 | -0.011 | 0.99 | -0.012 | 0.99 |
| Internal perceptual | 0.025 | 0.99 | 0.048 | 0.99 | -0.01 | 0.99 | -0.068 | 0.93 |
| Internal thoughts/emotions | 0.066 | 0.93 | 0.12 | 0.77 | -0.018 | 0.99 | -0.16 | 0.75 |
| Vividness rating | -0.11 | 0.80 | -0.062 | 0.97 | -0.15 | 0.75 | -0.048 | 0.99 |
| **Future thinking** |  |  |  |  |  |  |  |  |
| Experiential index | -0.039 | 0.87 | -0.031 | 0.90 | -0.038 | 0.87 | 0.005 | 0.98 |
| Spatial references | -0.052 | 0.81 | -0.076 | 0.60 | -0.01 | 0.97 | 0.083 | 0.55 |
| Entities present | -0.015 | 0.97 | -0.021 | 0.94 | -0.005 | 0.98 | 0.021 | 0.94 |
| Sensory descriptions | 0.021 | 0.94 | 0.012 | 0.97 | 0.027 | 0.93 | 0.007 | 0.98 |
| Thoughts/emotions/actions | -0.042 | 0.87 | -0.003 | 0.98 | -0.079 | 0.59 | -0.063 | 0.70 |
| Spatial coherence index | -0.016 | 0.97 | 0.00 | 1.00 | -0.032 | 0.90 | -0.026 | 0.93 |
| **Navigation** |  |  |  |  |  |  |  |  |
| Overall navigation score | -0.012 | 0.99 | -0.018 | 0.99 | -0.001 | 0.99 | 0.022 | 0.99 |
| Movie clip recognition | -0.13 | 0.91 | -0.10 | 0.91 | -0.14 | 0.91 | 0.012 | 0.99 |
| Scene recognition | -0.064 | 0.96 | -0.072 | 0.96 | -0.039 | 0.99 | 0.055 | 0.96 |
| Proximity judgements | 0.12 | 0.91 | 0.091 | 0.91 | 0.12 | 0.91 | -0.012 | 0.99 |
| Route knowledge | -0.015 | 0.99 | -0.05 | 0.99 | 0.032 | 0.99 | 0.087 | 0.92 |
| Sketch map | -0.009 | 0.99 | -0.01 | 0.99 | -0.005 | 0.99 | 0.009 | 0.99 |

P values are Benjamini-Hochberg false discovery rate corrected at $p < 0.05$ .

**Table S8.** Partial correlations between task performance and hippocampal grey matter <u>R_1_</u> with age, gender, total intracranial volume and MRI scanner as covariates.

| Performance variable | Whole hippocampus |  | Anterior hippocampus |  | Posterior hippocampus |  | Posterior/Anterior hippocampus ratio |  |
| --- | --- | --- | --- | --- | --- | --- | --- | --- |
|  | r | p | r | p | r | p | r | p |
| **Scene construction** |  |  |  |  |  |  |  |  |
| Experiential index | 0.092 | 0.59 | 0.11 | 0.59 | 0.065 | 0.67 | -0.082 | 0.59 |
| Spatial references | 0.11 | 0.59 | 0.13 | 0.55 | 0.087 | 0.59 | -0.07 | 0.63 |
| Entities present | 0.12 | 0.59 | 0.13 | 0.59 | 0.10 | 0.59 | -0.044 | 0.73 |
| Sensory descriptions | 0.061 | 0.67 | 0.06 | 0.67 | 0.055 | 0.71 | -0.018 | 0.87 |
| Thoughts/emotions/actions | 0.16 | 0.55 | 0.15 | 0.55 | 0.15 | 0.55 | -0.001 | 0.99 |
| Spatial coherence index | -0.039 | 0.77 | -0.035 | 0.77 | -0.035 | 0.77 | 0.013 | 0.91 |
| **Autobiographical interview** |  |  |  |  |  |  |  |  |
| Internal details | 0.13 | 0.77 | 0.091 | 0.85 | 0.15 | 0.75 | 0.08 | 0.88 |
| Internal events | 0.14 | 0.75 | 0.088 | 0.88 | 0.16 | 0.75 | 0.11 | 0.85 |
| Internal time | 0.072 | 0.93 | 0.077 | 0.88 | 0.061 | 0.97 | -0.036 | 0.99 |
| Internal place | -0.003 | 0.99 | -0.01 | 0.99 | 0.003 | 0.99 | 0.013 | 0.99 |
| Internal perceptual | 0.12 | 0.77 | 0.11 | 0.85 | 0.12 | 0.77 | 0.013 | 0.99 |
| Internal thoughts/emotions | -0.001 | 1.00 | -0.03 | 0.99 | 0.023 | 0.99 | 0.084 | 0.88 |
| Vividness rating | -0.034 | 0.99 | -0.038 | 0.99 | -0.027 | 0.99 | 0.008 | 0.99 |
| **Future thinking** |  |  |  |  |  |  |  |  |
| Experiential index | 0.13 | 0.31 | 0.12 | 0.34 | 0.13 | 0.31 | 0.006 | 0.98 |
| Spatial references | 0.18 | 0.31 | 0.17 | 0.31 | 0.17 | 0.31 | -0.01 | 0.97 |
| Entities present | 0.16 | 0.31 | 0.16 | 0.31 | 0.14 | 0.31 | -0.033 | 0.90 |
| Sensory descriptions | 0.11 | 0.38 | 0.11 | 0.38 | 0.098 | 0.40 | -0.014 | 0.97 |
| Thoughts/emotions/actions | 0.11 | 0.38 | 0.063 | 0.70 | 0.14 | 0.31 | 0.11 | 0.38 |
| Spatial coherence index | -0.069 | 0.70 | -0.064 | 0.70 | -0.067 | 0.70 | 0.005 | 0.98 |
| **Navigation** |  |  |  |  |  |  |  |  |
| Overall navigation score | -0.014 | 0.99 | 0.002 | 0.99 | -0.026 | 0.99 | -0.045 | 0.99 |
| Movie clip recognition | 0.021 | 0.99 | 0.056 | 0.96 | -0.011 | 0.99 | -0.11 | 0.91 |
| Scene recognition | 0.02 | 0.99 | 0.038 | 0.99 | 0.003 | 0.99 | -0.055 | 0.96 |
| Proximity judgements | -0.07 | 0.96 | -0.052 | 0.98 | -0.078 | 0.96 | -0.033 | 0.99 |
| Route knowledge | 0.004 | 0.99 | 0.038 | 0.99 | -0.025 | 0.99 | -0.11 | 0.91 |
| Sketch map | -0.015 | 0.99 | -0.004 | 0.99 | -0.023 | 0.99 | -0.029 | 0.99 |

P values are Benjamini-Hochberg false discovery rate corrected at $p < 0.05$ .

**Table S9.** Partial correlations between task performance and hippocampal grey matter <u>R_2_*</u> with age, gender, total intracranial volume and MRI scanner as covariates.

| Performance variable | Whole hippocampus |  | Anterior hippocampus |  | Posterior hippocampus |  | Posterior/Anterior hippocampus ratio |  |
| --- | --- | --- | --- | --- | --- | --- | --- | --- |
|  | r | p | r | p | r | p | r | p |
| **Scene construction** |  |  |  |  |  |  |  |  |
| Experiential index | 0.082 | 0.59 | 0.059 | 0.67 | 0.086 | 0.59 | 0.004 | 0.98 |
| Spatial references | 0.12 | 0.59 | 0.076 | 0.60 | 0.13 | 0.55 | 0.026 | 0.83 |
| Entities present | 0.10 | 0.59 | 0.047 | 0.73 | 0.13 | 0.55 | 0.08 | 0.59 |
| Sensory descriptions | 0.05 | 0.73 | 0.024 | 0.83 | 0.065 | 0.67 | 0.047 | 0.73 |
| Thoughts/emotions/actions | 0.081 | 0.59 | -0.005 | 0.98 | 0.15 | 0.55 | 0.13 | 0.55 |
| Spatial coherence index | -0.018 | 0.87 | 0.032 | 0.81 | -0.064 | 0.67 | -0.08 | 0.59 |
| **Autobiographical interview** |  |  |  |  |  |  |  |  |
| Internal details | 0.029 | 0.99 | 0.002 | 0.99 | 0.049 | 0.99 | 0.034 | 0.99 |
| Internal events | 0.028 | 0.99 | -0.004 | 0.99 | 0.055 | 0.99 | 0.052 | 0.99 |
| Internal time | -0.048 | 0.99 | -0.079 | 0.88 | -0.005 | 0.99 | 0.079 | 0.88 |
| Internal place | -0.042 | 0.99 | -0.043 | 0.99 | -0.031 | 0.99 | 0.019 | 0.99 |
| Internal perceptual | 0.066 | 0.93 | 0.057 | 0.99 | 0.059 | 0.98 | -0.019 | 0.99 |
| Internal thoughts/emotions | -0.027 | 0.99 | -0.039 | 0.99 | -0.01 | 0.99 | 0.021 | 0.99 |
| Vividness rating | -0.04 | 0.99 | -0.041 | 0.99 | -0.029 | 0.99 | 0.025 | 0.99 |
| **Future thinking** |  |  |  |  |  |  |  |  |
| Experiential index | 0.12 | 0.35 | 0.11 | 0.37 | 0.099 | 0.40 | -0.032 | 0.90 |
| Spatial references | 0.15 | 0.31 | 0.13 | 0.31 | 0.14 | 0.31 | -0.014 | 0.97 |
| Entities present | 0.13 | 0.31 | 0.13 | 0.31 | 0.095 | 0.44 | -0.049 | 0.81 |
| Sensory descriptions | 0.11 | 0.37 | 0.11 | 0.37 | 0.084 | 0.54 | -0.025 | 0.94 |
| Thoughts/emotions/actions | 0.01 | 0.97 | -0.011 | 0.97 | 0.03 | 0.92 | 0.023 | 0.94 |
| Spatial coherence index | 0.078 | 0.59 | 0.10 | 0.40 | 0.038 | 0.87 | -0.061 | 0.70 |
| **Navigation** |  |  |  |  |  |  |  |  |
| Overall navigation score | 0.065 | 0.96 | 0.018 | 0.99 | 0.097 | 0.91 | 0.095 | 0.91 |
| Movie clip recognition | 0.01 | 0.99 | 0.023 | 0.99 | -0.006 | 0.99 | -0.015 | 0.99 |
| Scene recognition | 0.099 | 0.91 | 0.067 | 0.96 | 0.11 | 0.91 | 0.04 | 0.99 |
| Proximity judgements | 0.017 | 0.99 | 0.032 | 0.99 | -0.002 | 0.99 | -0.013 | 0.99 |
| Route knowledge | 0.016 | 0.99 | -0.036 | 0.99 | 0.065 | 0.96 | 0.11 | 0.91 |
| Sketch map | 0.065 | 0.96 | 0.02 | 0.99 | 0.096 | 0.91 | 0.092 | 0.91 |

P values are Benjamini-Hochberg false discovery rate corrected at $p < 0.05$ .

**Table S10.** Partial correlations between task performance and hippocampal grey matter <u>MT saturation</u> in the <u>male</u> participants with age, total intracranial volume and MRI scanner as covariates.

| Performance variable | Whole hippocampus |  | Anterior hippocampus |  | Posterior hippocampus |  | Posterior/Anterior hippocampus ratio |  |
| --- | --- | --- | --- | --- | --- | --- | --- | --- |
|  | r | p | r | p | r | p | r | p |
| **Scene construction** |  |  |  |  |  |  |  |  |
| Experiential index | -0.18 | 0.70 | -0.17 | 0.70 | -0.16 | 0.70 | 0.072 | 0.83 |
| Spatial references | -0.075 | 0.83 | -0.094 | 0.76 | -0.041 | 0.86 | 0.095 | 0.76 |
| Entities present | -0.072 | 0.84 | -0.089 | 0.79 | -0.039 | 0.86 | 0.11 | 0.76 |
| Sensory descriptions | -0.14 | 0.70 | -0.16 | 0.70 | -0.085 | 0.81 | 0.16 | 0.70 |
| Thoughts/emotions/actions | -0.069 | 0.84 | -0.097 | 0.76 | -0.026 | 0.87 | 0.10 | 0.76 |
| Spatial coherence index | -0.21 | 0.70 | -0.12 | 0.70 | -0.26 | 0.67 | -0.089 | 0.79 |
| **Autobiographical interview** |  |  |  |  |  |  |  |  |
| Internal details | -0.017 | 1.00 | -0.008 | 1.00 | -0.022 | 1.00 | -0.018 | 1.00 |
| Internal events | -0.018 | 1.00 | -0.011 | 1.00 | -0.022 | 1.00 | -0.008 | 1.00 |
| Internal time | 0.10 | 0.79 | 0.16 | 0.71 | 0.027 | 1.00 | -0.20 | 0.71 |
| Internal place | -0.064 | 0.94 | -0.052 | 0.98 | -0.065 | 0.94 | -0.007 | 1.00 |
| Internal perceptual | 0.043 | 0.98 | 0.007 | 1.00 | 0.073 | 0.94 | 0.066 | 0.94 |
| Internal thoughts/emotions | -0.13 | 0.73 | -0.067 | 0.94 | -0.16 | 0.71 | -0.082 | 0.90 |
| Vividness rating | -0.24 | 0.71 | -0.21 | 0.71 | -0.23 | 0.71 | 0.052 | 0.98 |
| **Future thinking** |  |  |  |  |  |  |  |  |
| Experiential index | -0.22 | 0.35 | -0.18 | 0.35 | -0.21 | 0.35 | 0.045 | 0.92 |
| Spatial references | -0.19 | 0.35 | -0.16 | 0.35 | -0.20 | 0.35 | 0.029 | 0.92 |
| Entities present | -0.076 | 0.84 | -0.055 | 0.87 | -0.084 | 0.78 | 0.02 | 0.93 |
| Sensory descriptions | -0.054 | 0.87 | -0.071 | 0.85 | -0.026 | 0.92 | 0.12 | 0.52 |
| Thoughts/emotions/actions | -0.15 | 0.35 | -0.14 | 0.42 | -0.14 | 0.41 | 0.036 | 0.92 |
| Spatial coherence index | 0.20 | 0.35 | -0.14 | 0.42 | -0.23 | 0.35 | -0.029 | 0.92 |
| **Navigation** |  |  |  |  |  |  |  |  |
| Overall navigation score | -0.092 | 0.95 | -0.057 | 0.95 | -0.11 | 0.95 | -0.039 | 0.95 |
| Movie clip recognition | 0.00 | 1.00 | 0.037 | 0.95 | -0.039 | 0.95 | -0.079 | 0.95 |
| Scene recognition | 0.11 | 0.95 | 0.11 | 0.95 | 0.087 | 0.95 | -0.066 | 0.95 |
| Proximity judgements | -0.014 | 0.95 | 0.002 | 1.00 | -0.029 | 0.95 | -0.018 | 0.95 |
| Route knowledge | 0.017 | 0.95 | -0.021 | 0.95 | 0.054 | 0.95 | 0.083 | 0.95 |
| Sketch map | -0.12 | 0.95 | -0.071 | 0.95 | -0.14 | 0.95 | -0.052 | 0.95 |

P values are Benjamini-Hochberg false discovery rate corrected at $p < 0.05$ .

**Table S11.** Partial correlations between task performance and hippocampal grey matter <u>PD</u> in the <u>male</u> participants with age, total intracranial volume and MRI scanner as covariates.

| Performance variable | Whole hippocampus |  | Anterior hippocampus |  | Posterior hippocampus |  | Posterior/Anterior hippocampus ratio |  |
| --- | --- | --- | --- | --- | --- | --- | --- | --- |
|  | r | p | r | p | r | p | r | p |
| **Scene construction** |  |  |  |  |  |  |  |  |
| Experiential index | 0.073 | 0.83 | 0.034 | 0.86 | 0.098 | 0.76 | 0.038 | 0.86 |
| Spatial references | 0.045 | 0.85 | -0.055 | 0.85 | 0.15 | 0.70 | 0.18 | 0.70 |
| Entities present | 0.005 | 0.97 | -0.018 | 0.92 | 0.031 | 0.86 | 0.046 | 0.85 |
| Sensory descriptions | -0.046 | 0.85 | -0.046 | 0.85 | -0.032 | 0.86 | 0.027 | 0.87 |
| Thoughts/emotions/actions | 0.13 | 0.70 | 0.099 | 0.76 | 0.12 | 0.70 | -0.019 | 0.92 |
| Spatial coherence index | -0.052 | 0.85 | -0.03 | 0.86 | -0.062 | 0.85 | -0.015 | 0.93 |
| **Autobiographical interview** |  |  |  |  |  |  |  |  |
| Internal details | 0.086 | 0.88 | 0.14 | 0.71 | -0.007 | 1.00 | -0.16 | 0.71 |
| Internal events | 0.088 | 0.86 | 0.14 | 0.71 | 0.00 | 1.00 | -0.16 | 0.71 |
| Internal time | -0.047 | 0.98 | 0.016 | 1.00 | -0.11 | 0.79 | -0.10 | 0.80 |
| Internal place | 0.063 | 0.94 | 0.044 | 0.98 | 0.067 | 0.94 | 0.001 | 1.00 |
| Internal perceptual | 0.065 | 0.94 | 0.089 | 0.86 | 0.017 | 1.00 | -0.087 | 0.87 |
| Internal thoughts/emotions | 0.05 | 0.98 | 0.11 | 0.79 | -0.032 | 1.00 | -0.14 | 0.71 |
| Vividness rating | -0.13 | 0.73 | -0.077 | 0.92 | -0.16 | 0.71 | -0.04 | 1.00 |
| **Future thinking** |  |  |  |  |  |  |  |  |
| Experiential index | 0.033 | 0.92 | 0.056 | 0.87 | -0.004 | 0.97 | -0.066 | 0.86 |
| Spatial references | -0.028 | 0.92 | -0.052 | 0.89 | 0.01 | 0.94 | 0.063 | 0.86 |
| Entities present | 0.063 | 0.86 | 0.037 | 0.92 | 0.074 | 0.85 | 0.017 | 0.93 |
| Sensory descriptions | -0.038 | 0.92 | -0.009 | 0.94 | -0.06 | 0.86 | -0.036 | 0.92 |
| Thoughts/emotions/actions | 0.07 | 0.85 | 0.13 | 0.46 | -0.022 | 0.93 | -0.16 | 0.35 |
| Spatial coherence index | 0.024 | 0.93 | 0.01 | 0.94 | 0.033 | 0.92 | 0.014 | 0.93 |
| **Navigation** |  |  |  |  |  |  |  |  |
| Overall navigation score | 0.08 | 0.95 | 0.11 | 0.95 | 0.02 | 0.95 | -0.10 | 0.95 |
| Movie clip recognition | -0.17 | 0.95 | -0.12 | 0.95 | -0.19 | 0.95 | -0.012 | 0.95 |
| Scene recognition | 0.013 | 0.95 | 0.052 | 0.95 | -0.039 | 0.95 | -0.086 | 0.95 |
| Proximity judgements | 0.18 | 0.95 | 0.12 | 0.95 | 0.20 | 0.95 | 0.024 | 0.95 |
| Route knowledge | 0.081 | 0.95 | 0.051 | 0.95 | 0.093 | 0.95 | 0.016 | 0.95 |
| Sketch map | 0.077 | 0.95 | 0.11 | 0.95 | 0.008 | 0.97 | -0.12 | 0.95 |

P values are Benjamini-Hochberg false discovery rate corrected at $p < 0.05$ .

**Table S12.** Partial correlations between task performance and hippocampal grey matter <u>R_1_</u> in the <u>male</u> participants with age, total intracranial volume and MRI scanner as covariates.

| Performance variable | Whole hippocampus |  | Anterior hippocampus |  | Posterior hippocampus |  | Posterior/Anterior hippocampus ratio |  |
| --- | --- | --- | --- | --- | --- | --- | --- | --- |
|  | r | p | r | p | r | p | r | p |
| **Scene construction** |  |  |  |  |  |  |  |  |
| Experiential index | 0.055 | 0.85 | 0.072 | 0.83 | 0.034 | 0.86 | -0.059 | 0.85 |
| Spatial references | 0.083 | 0.82 | 0.13 | 0.70 | 0.039 | 0.86 | -0.13 | 0.70 |
| Entities present | 0.08 | 0.83 | 0.099 | 0.76 | 0.055 | 0.85 | -0.066 | 0.84 |
| Sensory descriptions | 0.13 | 0.70 | 0.13 | 0.70 | 0.12 | 0.70 | -0.03 | 0.86 |
| Thoughts/emotions/actions | 0.15 | 0.70 | 0.094 | 0.76 | 0.17 | 0.70 | 0.11 | 0.73 |
| Spatial coherence index | -0.053 | 0.85 | -0.051 | 0.85 | -0.049 | 0.85 | 0.014 | 0.93 |
| **Autobiographical interview** |  |  |  |  |  |  |  |  |
| Internal details | 0.18 | 0.71 | 0.14 | 0.71 | 0.19 | 0.71 | 0.06 | 0.95 |
| Internal events | 0.17 | 0.71 | 0.14 | 0.73 | 0.18 | 0.71 | 0.056 | 0.98 |
| Internal time | 0.15 | 0.71 | 0.12 | 0.75 | 0.15 | 0.71 | 0.044 | 0.98 |
| Internal place | 0.11 | 0.79 | 0.12 | 0.75 | 0.093 | 0.85 | -0.051 | 0.98 |
| Internal perceptual | 0.18 | 0.71 | 0.16 | 0.71 | 0.17 | 0.71 | 0.00 | 1.00 |
| Internal thoughts/emotions | -0.022 | 1.00 | -0.065 | 0.94 | 0.016 | 1.00 | 0.12 | 0.73 |
| Vividness rating | -0.002 | 1.00 | -0.017 | 1.00 | 0.01 | 1.00 | 0.026 | 1.00 |
| **Future thinking** |  |  |  |  |  |  |  |  |
| Experiential index | 0.16 | 0.35 | 0.12 | 0.48 | 0.17 | 0.35 | 0.063 | 0.86 |
| Spatial references | 0.17 | 0.35 | 0.15 | 0.37 | 0.17 | 0.35 | 0.024 | 0.93 |
| Entities present | 0.14 | 0.42 | 0.16 | 0.35 | 0.11 | 0.57 | -0.072 | 0.85 |
| Sensory descriptions | 0.19 | 0.35 | 0.17 | 0.35 | 0.19 | 0.35 | 0.028 | 0.92 |
| Thoughts/emotions/actions | 0.11 | 0.54 | 0.018 | 0.93 | 0.18 | 0.35 | 0.23 | 0.35 |
| Spatial coherence index | -0.051 | 0.89 | -0.038 | 0.92 | -0.056 | 0.87 | -0.015 | 0.93 |
| **Navigation** |  |  |  |  |  |  |  |  |
| Overall navigation score | -0.034 | 0.95 | -0.043 | 0.95 | -0.022 | 0.95 | 0.029 | 0.95 |
| Movie clip recognition | -0.012 | 0.95 | 0.017 | 0.95 | -0.035 | 0.95 | -0.075 | 0.95 |
| Scene recognition | 0.028 | 0.95 | 0.017 | 0.95 | 0.035 | 0.95 | 0.027 | 0.95 |
| Proximity judgements | 0.033 | 0.95 | 0.042 | 0.95 | 0.022 | 0.95 | -0.02 | 0.95 |
| Route knowledge | 0.033 | 0.95 | 0.049 | 0.95 | 0.017 | 0.95 | -0.052 | 0.95 |
| Sketch map | -0.047 | 0.95 | -0.061 | 0.95 | -0.031 | 0.95 | 0.043 | 0.95 |

P values are Benjamini-Hochberg false discovery rate corrected at $p < 0.05$ .

**Table S13.**
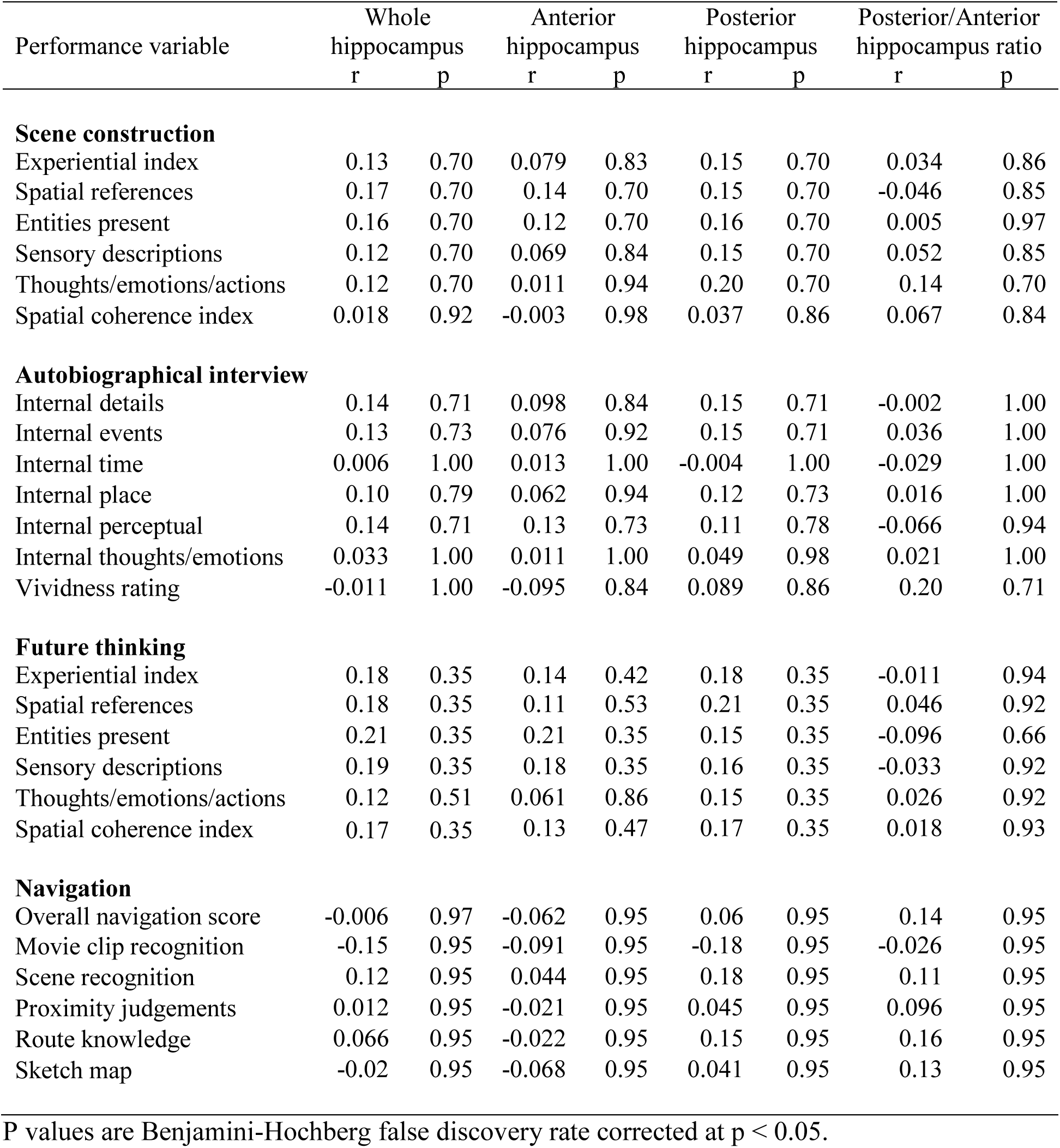
Partial correlations between task performance and hippocampal grey matter <u>R_2_*</u> in the <u>male</u> participants with age, total intracranial volume and MRI scanner as covariates.

| Performance variable | Whole hippocampus |  | Anterior hippocampus |  | Posterior hippocampus |  | Posterior/Anterior hippocampus ratio |  |
| --- | --- | --- | --- | --- | --- | --- | --- | --- |
|  | r | p | r | p | r | p | r | p |
| **Scene construction** |  |  |  |  |  |  |  |  |
| Experiential index | 0.13 | 0.70 | 0.079 | 0.83 | 0.15 | 0.70 | 0.034 | 0.86 |
| Spatial references | 0.17 | 0.70 | 0.14 | 0.70 | 0.15 | 0.70 | -0.046 | 0.85 |
| Entities present | 0.16 | 0.70 | 0.12 | 0.70 | 0.16 | 0.70 | 0.005 | 0.97 |
| Sensory descriptions | 0.12 | 0.70 | 0.069 | 0.84 | 0.15 | 0.70 | 0.052 | 0.85 |
| Thoughts/emotions/actions | 0.12 | 0.70 | 0.011 | 0.94 | 0.20 | 0.70 | 0.14 | 0.70 |
| Spatial coherence index | 0.018 | 0.92 | -0.003 | 0.98 | 0.037 | 0.86 | 0.067 | 0.84 |
| **Autobiographical interview** |  |  |  |  |  |  |  |  |
| Internal details | 0.14 | 0.71 | 0.098 | 0.84 | 0.15 | 0.71 | -0.002 | 1.00 |
| Internal events | 0.13 | 0.73 | 0.076 | 0.92 | 0.15 | 0.71 | 0.036 | 1.00 |
| Internal time | 0.006 | 1.00 | 0.013 | 1.00 | -0.004 | 1.00 | -0.029 | 1.00 |
| Internal place | 0.10 | 0.79 | 0.062 | 0.94 | 0.12 | 0.73 | 0.016 | 1.00 |
| Internal perceptual | 0.14 | 0.71 | 0.13 | 0.73 | 0.11 | 0.78 | -0.066 | 0.94 |
| Internal thoughts/emotions | 0.033 | 1.00 | 0.011 | 1.00 | 0.049 | 0.98 | 0.021 | 1.00 |
| Vividness rating | -0.011 | 1.00 | -0.095 | 0.84 | 0.089 | 0.86 | 0.20 | 0.71 |
| **Future thinking** |  |  |  |  |  |  |  |  |
| Experiential index | 0.18 | 0.35 | 0.14 | 0.42 | 0.18 | 0.35 | -0.011 | 0.94 |
| Spatial references | 0.18 | 0.35 | 0.11 | 0.53 | 0.21 | 0.35 | 0.046 | 0.92 |
| Entities present | 0.21 | 0.35 | 0.21 | 0.35 | 0.15 | 0.35 | -0.096 | 0.66 |
| Sensory descriptions | 0.19 | 0.35 | 0.18 | 0.35 | 0.16 | 0.35 | -0.033 | 0.92 |
| Thoughts/emotions/actions | 0.12 | 0.51 | 0.061 | 0.86 | 0.15 | 0.35 | 0.026 | 0.92 |
| Spatial coherence index | 0.17 | 0.35 | 0.13 | 0.47 | 0.17 | 0.35 | 0.018 | 0.93 |
| **Navigation** |  |  |  |  |  |  |  |  |
| Overall navigation score | -0.006 | 0.97 | -0.062 | 0.95 | 0.06 | 0.95 | 0.14 | 0.95 |
| Movie clip recognition | -0.15 | 0.95 | -0.091 | 0.95 | -0.18 | 0.95 | -0.026 | 0.95 |
| Scene recognition | 0.12 | 0.95 | 0.044 | 0.95 | 0.18 | 0.95 | 0.11 | 0.95 |
| Proximity judgements | 0.012 | 0.95 | -0.021 | 0.95 | 0.045 | 0.95 | 0.096 | 0.95 |
| Route knowledge | 0.066 | 0.95 | -0.022 | 0.95 | 0.15 | 0.95 | 0.16 | 0.95 |
| Sketch map | -0.02 | 0.95 | -0.068 | 0.95 | 0.041 | 0.95 | 0.13 | 0.95 |

P values are Benjamini-Hochberg false discovery rate corrected at $p < 0.05$ .

**Table S14.** Partial correlations between tasks performance and hippocampal grey matter <u>MT saturation</u> in the <u>female</u> participants with age, total intracranial volume and MRI scanner as covariates.

| Performance variable | Whole hippocampus |  | Anterior hippocampus |  | Posterior hippocampus |  | Posterior/Anterior hippocampus ratio |  |
| --- | --- | --- | --- | --- | --- | --- | --- | --- |
|  | r | p | r | p | r | p | r | p |
| **Scene construction** |  |  |  |  |  |  |  |  |
| Experiential index | -0.002 | 1.00 | -0.005 | 1.00 | 0.001 | 1.00 | -0.001 | 1.00 |
| Spatial references | -0.076 | 0.95 | -0.070 | 0.96 | -0.074 | 0.95 | 0.004 | 1.00 |
| Entities present | -0.048 | 0.96 | -0.059 | 0.96 | -0.034 | 0.96 | 0.035 | 0.96 |
| Sensory descriptions | -0.084 | 0.94 | -0.093 | 0.88 | -0.067 | 0.96 | 0.034 | 0.96 |
| Thoughts/emotions/actions | -0.002 | 1.00 | -0.034 | 0.96 | 0.027 | 0.96 | 0.095 | 0.88 |
| Spatial coherence index | 0.065 | 0.96 | 0.081 | 0.95 | 0.044 | 0.96 | -0.089 | 0.91 |
| **Autobiographical interview** |  |  |  |  |  |  |  |  |
| Internal details | 0.010 | 1.00 | 0.012 | 1.00 | 0.007 | 1.00 | -0.023 | 1.00 |
| Internal events | 0.021 | 1.00 | 0.022 | 1.00 | 0.018 | 1.00 | -0.020 | 1.00 |
| Internal time | 0.020 | 1.00 | 0.051 | 1.00 | -0.009 | 1.00 | -0.099 | 1.00 |
| Internal place | -0.078 | 1.00 | -0.050 | 1.00 | -0.095 | 1.00 | -0.069 | 1.00 |
| Internal perceptual | -0.033 | 1.00 | -0.030 | 1.00 | -0.032 | 1.00 | -0.009 | 1.00 |
| Internal thoughts/emotions | 0.084 | 1.00 | 0.062 | 1.00 | 0.096 | 1.00 | 0.033 | 1.00 |
| Vividness rating | -0.011 | 1.00 | 0.000 | 1.00 | -0.019 | 1.00 | -0.044 | 1.00 |
| **Future thinking** |  |  |  |  |  |  |  |  |
| Experiential index | -0.035 | 0.97 | -0.054 | 0.89 | -0.015 | 1.00 | 0.054 | 0.89 |
| Spatial references | -0.11 | 0.81 | -0.123 | 0.81 | -0.094 | 0.81 | 0.062 | 0.89 |
| Entities present | 0.001 | 1.00 | -0.009 | 1.00 | 0.010 | 1.00 | 0.018 | 1.00 |
| Sensory descriptions | -0.13 | 0.81 | -0.146 | 0.81 | -0.104 | 0.81 | 0.076 | 0.83 |
| Thoughts/emotions/actions | 0.13 | 0.81 | 0.087 | 0.81 | 0.151 | 0.81 | 0.059 | 0.89 |
| Spatial coherence index | 0.019 | 1.00 | 0.047 | 0.93 | -0.009 | 1.00 | -0.104 | 0.81 |
| **Navigation** |  |  |  |  |  |  |  |  |
| Overall navigation score | -0.10 | 0.71 | -0.059 | 0.88 | -0.13 | 0.65 | -0.081 | 0.80 |
| Movie clip recognition | 0.087 | 0.79 | 0.091 | 0.76 | 0.075 | 0.82 | -0.029 | 0.91 |
| Scene recognition | 0.047 | 0.91 | 0.042 | 0.91 | 0.047 | 0.91 | -0.001 | 0.99 |
| Proximity judgements | -0.13 | 0.65 | -0.12 | 0.71 | -0.13 | 0.65 | 0.008 | 0.97 |
| Route knowledge | 0.013 | 0.96 | 0.071 | 0.82 | -0.040 | 0.91 | -0.18 | 0.65 |
| Sketch map | -0.12 | 0.71 | -0.078 | 0.82 | -0.15 | 0.65 | -0.066 | 0.85 |

P values are Benjamini-Hochberg false discovery rate corrected at $p < 0.05$ .

**Table S15.** Partial correlations between tasks performance and hippocampal grey matter <u>PD</u> in the <u>female</u> participants with age, total intracranial volume and MRI scanner as covariates.

| Performance variable | Whole hippocampus |  | Anterior hippocampus |  | Posterior hippocampus |  | Posterior/Anterior hippocampus ratio |  |
| --- | --- | --- | --- | --- | --- | --- | --- | --- |
|  | r | p | r | p | r | p | r | p |
| **Scene construction** |  |  |  |  |  |  |  |  |
| Experiential index | -0.12 | 0.88 | -0.12 | 0.88 | -0.11 | 0.88 | 0.058 | 0.96 |
| Spatial references | -0.080 | 0.95 | -0.098 | 0.88 | -0.044 | 0.96 | 0.094 | 0.88 |
| Entities present | -0.16 | 0.77 | -0.14 | 0.77 | -0.15 | 0.77 | 0.060 | 0.96 |
| Sensory descriptions | 0.095 | 0.88 | 0.066 | 0.96 | 0.11 | 0.88 | 0.012 | 0.98 |
| Thoughts/emotions/actions | -0.16 | 0.77 | -0.14 | 0.77 | -0.16 | 0.77 | 0.049 | 0.96 |
| Spatial coherence index | -0.031 | 0.96 | 0.029 | 0.96 | -0.098 | 0.88 | -0.13 | 0.83 |
| **Autobiographical interview** |  |  |  |  |  |  |  |  |
| Internal details | 0.005 | 1.00 | 0.037 | 1.00 | -0.036 | 1.00 | -0.089 | 1.00 |
| Internal events | 0.013 | 1.00 | 0.051 | 1.00 | -0.035 | 1.00 | -0.11 | 1.00 |
| Internal time | -0.11 | 1.00 | -0.16 | 1.00 | -0.033 | 1.00 | 0.19 | 1.00 |
| Internal place | -0.074 | 1.00 | -0.077 | 1.00 | -0.057 | 1.00 | 0.048 | 1.00 |
| Internal perceptual | -0.008 | 1.00 | 0.003 | 1.00 | -0.021 | 1.00 | -0.026 | 1.00 |
| Internal thoughts/emotions | 0.084 | 1.00 | 0.14 | 1.00 | 0.002 | 1.00 | -0.19 | 1.00 |
| Vividness rating | -0.095 | 1.00 | -0.045 | 1.00 | -0.14 | 1.00 | -0.072 | 1.00 |
| **Future thinking** |  |  |  |  |  |  |  |  |
| Experiential index | -0.13 | 0.81 | -0.14 | 0.81 | -0.090 | 0.81 | 0.11 | 0.81 |
| Spatial references | -0.068 | 0.87 | -0.086 | 0.81 | -0.033 | 0.98 | 0.088 | 0.81 |
| Entities present | -0.12 | 0.81 | -0.11 | 0.81 | -0.11 | 0.81 | 0.046 | 0.93 |
| Sensory descriptions | 0.096 | 0.81 | 0.052 | 0.89 | 0.13 | 0.81 | 0.052 | 0.89 |
| Thoughts/emotions/actions | -0.15 | 0.81 | -0.13 | 0.81 | -0.14 | 0.81 | 0.040 | 0.94 |
| Spatial coherence index | -0.059 | 0.89 | -0.012 | 1.00 | -0.11 | 0.81 | -0.085 | 0.81 |
| **Navigation** |  |  |  |  |  |  |  |  |
| Overall navigation score | -0.10 | 0.71 | -0.16 | 0.65 | -0.022 | 0.92 | 0.19 | 0.65 |
| Movie clip recognition | -0.11 | 0.71 | -0.11 | 0.71 | -0.096 | 0.75 | 0.058 | 0.88 |
| Scene recognition | -0.13 | 0.65 | -0.18 | 0.65 | -0.044 | 0.91 | 0.21 | 0.65 |
| Proximity judgements | 0.049 | 0.91 | 0.042 | 0.91 | 0.050 | 0.91 | -0.011 | 0.96 |
| Route knowledge | -0.10 | 0.71 | -0.15 | 0.65 | -0.032 | 0.91 | 0.17 | 0.65 |
| Sketch map | -0.093 | 0.76 | -0.14 | 0.65 | -0.017 | 0.95 | 0.18 | 0.65 |

P values are Benjamini-Hochberg false discovery rate corrected at $p < 0.05$ .

**Table S16.** Partial correlations between tasks performance and hippocampal grey matter <u>R_1_</u> in the <u>female</u> participants with age, total intracranial volume and MRI scanner as covariates.

| Performance variable | Whole hippocampus |  | Anterior hippocampus |  | Posterior hippocampus |  | Posterior/Anterior hippocampus ratio |  |
| --- | --- | --- | --- | --- | --- | --- | --- | --- |
|  | r | p | r | p | r | p | r | p |
| **Scene construction** |  |  |  |  |  |  |  |  |
| Experiential index | 0.13 | 0.88 | 0.15 | 0.77 | 0.094 | 0.88 | -0.10 | 0.88 |
| Spatial references | 0.15 | 0.77 | 0.15 | 0.77 | 0.14 | 0.77 | -0.019 | 0.96 |
| Entities present | 0.16 | 0.77 | 0.15 | 0.77 | 0.14 | 0.77 | -0.024 | 0.96 |
| Sensory descriptions | -0.017 | 0.96 | -0.021 | 0.96 | -0.012 | 0.98 | 0.007 | 1.00 |
| Thoughts/emotions/actions | 0.18 | 0.77 | 0.20 | 0.77 | 0.145 | 0.77 | -0.094 | 0.88 |
| Spatial coherence index | -0.033 | 0.96 | -0.040 | 0.96 | -0.025 | 0.96 | 0.021 | 0.96 |
| **Autobiographical interview** |  |  |  |  |  |  |  |  |
| Internal details | 0.11 | 1.00 | 0.070 | 1.00 | 0.13 | 1.00 | 0.084 | 1.00 |
| Internal events | 0.11 | 1.00 | 0.045 | 1.00 | 0.15 | 1.00 | 0.16 | 1.00 |
| Internal time | -0.010 | 1.00 | 0.022 | 1.00 | -0.037 | 1.00 | -0.10 | 1.00 |
| Internal place | -0.022 | 1.00 | -0.027 | 1.00 | -0.016 | 1.00 | 0.014 | 1.00 |
| Internal perceptual | 0.10 | 1.00 | 0.093 | 1.00 | 0.10 | 1.00 | 0.001 | 1.00 |
| Internal thoughts/emotions | 0.032 | 1.00 | 0.019 | 1.00 | 0.041 | 1.00 | 0.033 | 1.00 |
| Vividness rating | -0.082 | 1.00 | -0.080 | 1.00 | -0.077 | 1.00 | 0.000 | 1.00 |
| **Future thinking** |  |  |  |  |  |  |  |  |
| Experiential index | 0.12 | 0.81 | 0.13 | 0.81 | 0.094 | 0.81 | -0.064 | 0.89 |
| Spatial references | 0.16 | 0.81 | 0.16 | 0.81 | 0.14 | 0.81 | -0.030 | 0.99 |
| Entities present | 0.21 | 0.81 | 0.20 | 0.81 | 0.21 | 0.81 | 0.006 | 1.00 |
| Sensory descriptions | -0.002 | 1.00 | 0.015 | 1.00 | -0.016 | 1.00 | -0.054 | 0.89 |
| Thoughts/emotions/actions | 0.13 | 0.81 | 0.12 | 0.81 | 0.12 | 0.81 | 0.006 | 1.00 |
| Spatial coherence index | -0.072 | 0.85 | -0.077 | 0.83 | -0.062 | 0.89 | 0.025 | 1.00 |
| **Navigation** |  |  |  |  |  |  |  |  |
| Overall navigation score | 0.027 | 0.92 | 0.071 | 0.82 | -0.012 | 0.96 | -0.14 | 0.65 |
| Movie clip recognition | 0.069 | 0.83 | 0.12 | 0.71 | 0.024 | 0.92 | -0.15 | 0.65 |
| Scene recognition | 0.034 | 0.91 | 0.076 | 0.82 | -0.004 | 0.99 | -0.13 | 0.65 |
| Proximity judgements | -0.13 | 0.65 | -0.099 | 0.73 | -0.15 | 0.65 | -0.081 | 0.80 |
| Route knowledge | -0.018 | 0.95 | 0.034 | 0.91 | -0.059 | 0.88 | -0.16 | 0.65 |
| Sketch map | 0.035 | 0.91 | 0.072 | 0.82 | 0.001 | 0.99 | -0.11 | 0.71 |

P values are Benjamini-Hochberg false discovery rate corrected at $p < 0.05$ .

**Table S17.** Partial correlations between tasks performance and hippocampal grey matter <u>R_2_*</u> in the <u>female</u> participants with age, total intracranial volume and MRI scanner as covariates.

| Performance variable | Whole hippocampus |  | Anterior hippocampus |  | Posterior hippocampus |  | Posterior/Anterior hippocampus ratio |  |
| --- | --- | --- | --- | --- | --- | --- | --- | --- |
|  | r | p | r | p | r | p | r | p |
| **Scene construction** |  |  |  |  |  |  |  |  |
| Experiential index | 0.026 | 0.96 | 0.035 | 0.96 | 0.013 | 0.98 | -0.047 | 0.96 |
| Spatial references | 0.075 | 0.95 | 0.025 | 0.96 | 0.11 | 0.88 | 0.074 | 0.95 |
| Entities present | 0.050 | 0.96 | -0.018 | 0.96 | 0.10 | 0.88 | 0.14 | 0.77 |
| Sensory descriptions | -0.034 | 0.96 | -0.030 | 0.96 | -0.032 | 0.96 | 0.017 | 0.96 |
| Thoughts/emotions/actions | 0.041 | 0.96 | -0.018 | 0.96 | 0.088 | 0.91 | 0.096 | 0.88 |
| Spatial coherence index | -0.059 | 0.96 | 0.067 | 0.96 | -0.17 | 0.77 | -0.27 | 0.55 |
| **Autobiographical interview** |  |  |  |  |  |  |  |  |
| Internal details | -0.066 | 1.00 | -0.090 | 1.00 | -0.032 | 1.00 | 0.065 | 1.00 |
| Internal events | -0.062 | 1.00 | -0.092 | 1.00 | -0.022 | 1.00 | 0.085 | 1.00 |
| Internal time | -0.11 | 1.00 | -0.19 | 1.00 | -0.024 | 1.00 | 0.12 | 1.00 |
| Internal place | -0.12 | 1.00 | -0.11 | 1.00 | -0.11 | 1.00 | 0.025 | 1.00 |
| Internal perceptual | 0.019 | 1.00 | 0.015 | 1.00 | 0.020 | 1.00 | 0.001 | 1.00 |
| Internal thoughts/emotions | -0.087 | 1.00 | -0.084 | 1.00 | -0.073 | 1.00 | -0.006 | 1.00 |
| Vividness rating | -0.065 | 1.00 | 0.001 | 1.00 | -0.12 | 1.00 | -0.14 | 1.00 |
| **Future thinking** |  |  |  |  |  |  |  |  |
| Experiential index | 0.042 | 0.94 | 0.076 | 0.83 | 0.003 | 1.00 | -0.082 | 0.82 |
| Spatial references | 0.11 | 0.81 | 0.14 | 0.81 | 0.069 | 0.87 | -0.091 | 0.81 |
| Entities present | 0.041 | 0.94 | 0.029 | 0.99 | 0.043 | 0.94 | 0.023 | 1.00 |
| Sensory descriptions | 0.011 | 1.00 | 0.020 | 1.00 | 0.00 | 1.00 | -0.013 | 1.00 |
| Thoughts/emotions/actions | -0.10 | 0.81 | -0.087 | 0.81 | -0.094 | 0.81 | -0.003 | 1.00 |
| Spatial coherence index | -0.014 | 1.00 | 0.077 | 0.83 | -0.097 | 0.81 | -0.19 | 0.81 |
| **Navigation** |  |  |  |  |  |  |  |  |
| Overall navigation score | 0.13 | 0.65 | 0.12 | 0.71 | 0.12 | 0.71 | 0.006 | 0.98 |
| Movie clip recognition | 0.13 | 0.65 | 0.13 | 0.65 | 0.11 | 0.71 | -0.029 | 0.91 |
| Scene recognition | 0.085 | 0.79 | 0.091 | 0.76 | 0.064 | 0.85 | -0.029 | 0.91 |
| Proximity judgements | 0.039 | 0.91 | 0.11 | 0.71 | -0.033 | 0.91 | -0.15 | 0.65 |
| Route knowledge | -0.043 | 0.91 | -0.048 | 0.91 | -0.031 | 0.91 | 0.024 | 0.92 |
| Sketch map | 0.15 | 0.65 | 0.13 | 0.65 | 0.13 | 0.65 | 0.014 | 0.96 |

P values are Benjamini-Hochberg false discovery rate corrected at $p < 0.05$ .

**Table S18.** Details of the groups created for each task when dividing by median performance.

| Performance variable | Median performance | N (low performance) | N (high performance) |
| --- | --- | --- | --- |
| **Scene construction** |  |  |  |
| Experiential index | 41.2 | 109 | 108 |
| Spatial references | 3.29 | 112 | 105 |
| Entities present | 9.71 | 112 | 105 |
| Sensory descriptions | 12.14 | 112 | 105 |
| Thoughts/emotions/actions | 3.14 | 114 | 103 |
| Spatial coherence index | 3.0 | 112 | 105 |
| **Autobiographical interview** |  |  |  |
| Internal details | 23.2 | 110 | 107 |
| Internal events | 10.6 | 111 | 106 |
| Internal time | 1.4 | 128 | 89 |
| Internal place | 2.2 | 123 | 94 |
| Internal perceptual | 5.4 | 109 | 108 |
| Internal thoughts/emotions | 3.2 | 111 | 106 |
| Vividness rating | 4.6 | 114 | 103 |
| **Future thinking** |  |  |  |
| Experiential index | 39.8 | 111 | 106 |
| Spatial references | 2.3 | 123 | 94 |
| Entities present | 10.0 | 114 | 103 |
| Sensory descriptions | 8.67 | 117 | 100 |
| Thoughts/emotions/actions | 5.0 | 116 | 101 |
| Spatial coherence index | 2.67 | 115 | 102 |
| **Navigation** |  |  |  |
| Overall navigation score | 144.0 | 109 | 108 |
| Scene recognition | 30.0 | 146 | 71 |
| Proximity judgements | 8.0 | 159 | 58 |
| Route knowledge | 11.0 | 114 | 103 |
| Sketch map | 81 | 111 | 106 |

**Table S19.** Comparison of hippocampal grey matter <u>MT saturation</u> when dividing the sample into two groups determined by their <u>median performance</u> on each cognitive task, with age, gender, total intracranial volume and MRI scanner included as covariates.

| Performance variable | Whole hippocampus |  | Anterior hippocampus |  | Posterior hippocampus |  | Posterior/Anterior hippocampus ratio |  |
| --- | --- | --- | --- | --- | --- | --- | --- | --- |
|  | F | p | F | p | F | p | F | p |
| **Scene construction** |  |  |  |  |  |  |  |  |
| Experiential index | 0.03 | 0.96 | 0.08 | 0.92 | 0.00 | 0.99 | 0.26 | 0.88 |
| Spatial references | 0.91 | 0.75 | 0.86 | 0.75 | 0.71 | 0.75 | 0.36 | 0.88 |
| Entities present | 0.15 | 0.88 | 0.31 | 0.88 | 0.03 | 0.96 | 0.73 | 0.75 |
| Sensory descriptions | 2.51 | 0.52 | 4.19 | 0.42 | 0.84 | 0.75 | 4.22 | 0.42 |
| Thoughts/emotions/actions | 1.06 | 0.75 | 0.48 | 0.84 | 1.48 | 0.74 | 0.05 | 0.96 |
| Spatial coherence index | 0.40 | 0.88 | 0.14 | 0.88 | 2.33 | 0.56 | 6.13 | 0.42 |
| **Autobiographical interview** |  |  |  |  |  |  |  |  |
| Internal details | 0.14 | 0.95 | 0.10 | 0.95 | 0.14 | 0.95 | 0.03 | 0.96 |
| Internal events | 0.82 | 0.95 | 0.89 | 0.95 | 0.56 | 0.95 | 0.11 | 0.95 |
| Internal time | 0.00 | 1.00 | 0.33 | 0.95 | 0.31 | 0.95 | 2.67 | 0.95 |
| Internal place | 0.97 | 0.95 | 0.51 | 0.95 | 1.23 | 0.95 | 0.27 | 0.95 |
| Internal perceptual | 0.03 | 0.96 | 0.40 | 0.95 | 0.09 | 0.95 | 1.35 | 0.95 |
| Internal thoughts/emotions | 0.09 | 0.95 | 0.17 | 0.95 | 0.02 | 0.98 | 0.34 | 0.95 |
| Vividness ratings | 1.22 | 0.95 | 2.22 | 0.95 | 0.33 | 0.95 | 1.59 | 0.95 |
| **Future thinking** |  |  |  |  |  |  |  |  |
| Experiential index | 0.27 | 0.95 | 0.46 | 0.95 | 0.08 | 0.95 | 0.39 | 0.95 |
| Spatial references | 1.59 | 0.83 | 1.10 | 0.86 | 1.66 | 0.83 | 0.04 | 0.97 |
| Entities present | 0.68 | 0.95 | 0.63 | 0.95 | 0.54 | 0.95 | 0.03 | 0.97 |
| Sensory descriptions | 0.03 | 0.97 | 0.54 | 0.95 | 0.16 | 0.95 | 2.77 | 0.54 |
| Thoughts/emotions/actions | 3.11 | 0.54 | 2.95 | 0.54 | 2.43 | 0.58 | 0.26 | 0.95 |
| Spatial coherence index | 1.36 | 0.83 | 0.25 | 0.95 | 2.75 | 0.54 | 1.54 | 0.83 |
| **Navigation** |  |  |  |  |  |  |  |  |
| Overall navigation score | 6.28 | 0.37 | 4.75 | 0.40 | 6.08 | 0.37 | 0.07 | 0.96 |
| Scene recognition | 0.12 | 0.96 | 0.17 | 0.96 | 0.05 | 0.96 | 0.06 | 0.96 |
| Proximity judgements | 0.08 | 0.96 | 0.09 | 0.96 | 0.64 | 0.96 | 1.75 | 0.96 |
| Route knowledge | 0.04 | 0.96 | 0.07 | 0.96 | 0.01 | 0.98 | 0.07 | 0.96 |
| Sketch map | 5.25 | 0.37 | 2.92 | 0.79 | 6.44 | 0.37 | 0.26 | 0.96 |

P values are Benjamini-Hochberg false discovery rate corrected at $p < 0.05$ .

**Table S20.** Comparison of hippocampal grey matter <u>PD</u> when dividing the sample into two groups determined by their <u>median performance</u> on each cognitive task, with age, gender, total intracranial volume and MRI scanner included as covariates.

| Performance variable | Whole hippocampus |  | Anterior hippocampus |  | Posterior hippocampus |  | Posterior/Anterior hippocampus ratio |  |
| --- | --- | --- | --- | --- | --- | --- | --- | --- |
|  | F | p | F | p | F | p | F | p |
| **Scene construction** |  |  |  |  |  |  |  |  |
| Experiential index | 0.76 | 0.75 | 1.00 | 0.75 | 0.25 | 0.88 | 0.69 | 0.75 |
| Spatial references | 0.32 | 0.88 | 0.86 | 0.75 | 0.00 | 0.99 | 1.33 | 0.75 |
| Entities present | 1.75 | 0.72 | 2.00 | 0.66 | 0.78 | 0.75 | 1.04 | 0.75 |
| Sensory descriptions | 0.74 | 0.75 | 0.22 | 0.88 | 1.27 | 0.75 | 0.14 | 0.88 |
| Thoughts/emotions/actions | 0.04 | 0.96 | 0.27 | 0.88 | 0.05 | 0.96 | 0.66 | 0.76 |
| Spatial coherence index | 0.44 | 0.85 | 0.01 | 0.97 | 1.44 | 0.74 | 0.79 | 0.75 |
| **Autobiographical interview** |  |  |  |  |  |  |  |  |
| Internal details | 0.20 | 0.95 | 1.64 | 0.95 | 0.45 | 0.95 | 4.70 | 0.95 |
| Internal events | 0.26 | 0.95 | 0.80 | 0.95 | 0.01 | 0.98 | 1.45 | 0.95 |
| Internal time | 1.87 | 0.95 | 1.03 | 0.95 | 2.12 | 0.95 | 0.00 | 1.00 |
| Internal place | 0.77 | 0.95 | 0.80 | 0.95 | 0.41 | 0.95 | 0.33 | 0.95 |
| Internal perceptual | 0.22 | 0.95 | 0.30 | 0.95 | 0.06 | 0.95 | 0.23 | 0.95 |
| Internal thoughts/emotions | 0.10 | 0.95 | 1.08 | 0.95 | 0.43 | 0.95 | 3.31 | 0.95 |
| Vividness ratings | 0.49 | 0.95 | 0.06 | 0.95 | 1.18 | 0.95 | 0.41 | 0.95 |
| **Future thinking** |  |  |  |  |  |  |  |  |
| Experiential index | 0.44 | 0.95 | 0.60 | 0.95 | 0.14 | 0.95 | 0.34 | 0.95 |
| Spatial references | 1.21 | 0.83 | 1.78 | 0.80 | 0.29 | 0.95 | 1.41 | 0.83 |
| Entities present | 0.01 | 0.98 | 0.01 | 0.98 | 0.01 | 0.98 | 0.00 | 0.99 |
| Sensory descriptions | 0.02 | 0.98 | 0.06 | 0.95 | 0.00 | 1.00 | 0.10 | 0.95 |
| Thoughts/emotions/actions | 2.92 | 0.54 | 1.22 | 0.83 | 4.11 | 0.48 | 0.13 | 0.95 |
| Spatial coherence index | 0.28 | 0.95 | 0.26 | 0.95 | 0.18 | 0.95 | 0.08 | 0.95 |
| **Navigation** |  |  |  |  |  |  |  |  |
| Overall navigation score | 0.04 | 0.96 | 0.00 | 1.00 | 0.14 | 0.96 | 0.09 | 0.96 |
| Scene recognition | 0.12 | 0.96 | 0.00 | 0.98 | 0.38 | 0.96 | 0.20 | 0.96 |
| Proximity judgements | 1.99 | 0.96 | 2.40 | 0.89 | 0.80 | 0.96 | 1.25 | 0.96 |
| Route knowledge | 0.21 | 0.96 | 0.04 | 0.96 | 1.34 | 0.96 | 1.51 | 0.96 |
| Sketch map | 0.25 | 0.96 | 0.05 | 0.96 | 0.50 | 0.96 | 0.08 | 0.96 |

P values are Benjamini-Hochberg false discovery rate corrected at $p < 0.05$ .

**Table S21.** Comparison of hippocampal grey matter <u>R_1_</u> when dividing the sample into two groups determined by their <u>median performance</u> on each cognitive task, with age, gender, total intracranial volume and MRI scanner included as covariates.

| Performance variable | Whole hippocampus |  | Anterior hippocampus |  | Posterior hippocampus |  | Posterior/Anterior hippocampus ratio |  |
| --- | --- | --- | --- | --- | --- | --- | --- | --- |
|  | F | p | F | p | F | p | F | p |
| **Scene construction** |  |  |  |  |  |  |  |  |
| Experiential index | 2.75 | 0.50 | 3.60 | 0.47 | 1.68 | 0.72 | 0.92 | 0.75 |
| Spatial references | 4.13 | 0.42 | 4.83 | 0.42 | 2.88 | 0.50 | 0.64 | 0.76 |
| Entities present | 6.71 | 0.42 | 5.72 | 0.42 | 6.27 | 0.42 | 0.01 | 0.97 |
| Sensory descriptions | 0.09 | 0.92 | 0.18 | 0.88 | 0.03 | 0.96 | 0.18 | 0.88 |
| Thoughts/emotions/actions | 2.83 | 0.50 | 1.63 | 0.72 | 3.49 | 0.47 | 0.61 | 0.76 |
| Spatial coherence index | 0.24 | 0.88 | 0.20 | 0.88 | 0.23 | 0.88 | 0.00 | 0.99 |
| **Autobiographical interview** |  |  |  |  |  |  |  |  |
| Internal details | 7.60 | 0.41 | 3.72 | 0.95 | 10.23 | 0.20 | 2.75 | 0.95 |
| Internal events | 3.42 | 0.95 | 1.34 | 0.95 | 5.09 | 0.95 | 2.30 | 0.95 |
| Internal time | 3.73 | 0.95 | 2.47 | 0.95 | 4.20 | 0.95 | 0.31 | 0.95 |
| Internal place | 0.21 | 0.95 | 0.12 | 0.95 | 0.26 | 0.95 | 0.12 | 0.95 |
| Internal perceptual | 1.05 | 0.95 | 0.79 | 0.95 | 1.08 | 0.95 | 0.01 | 0.98 |
| Internal thoughts/emotions | 0.12 | 0.95 | 0.00 | 1.00 | 0.34 | 0.95 | 0.63 | 0.95 |
| Vividness ratings | 0.05 | 0.95 | 0.37 | 0.95 | 0.01 | 0.98 | 0.92 | 0.95 |
| **Future thinking** |  |  |  |  |  |  |  |  |
| Experiential index | 1.04 | 0.87 | 0.87 | 0.94 | 0.99 | 0.88 | 0.00 | 1.00 |
| Spatial references | 4.19 | 0.48 | 4.00 | 0.48 | 3.56 | 0.53 | 0.06 | 0.95 |
| Entities present | 5.17 | 0.48 | 4.14 | 0.48 | 5.08 | 0.48 | 0.06 | 0.95 |
| Sensory descriptions | 0.41 | 0.95 | 0.26 | 0.95 | 0.47 | 0.95 | 0.06 | 0.95 |
| Thoughts/emotions/actions | 3.89 | 0.48 | 1.47 | 0.83 | 5.90 | 0.48 | 2.93 | 0.54 |
| Spatial coherence index | 0.24 | 0.95 | 0.11 | 0.95 | 0.34 | 0.95 | 0.08 | 0.95 |
| **Navigation** |  |  |  |  |  |  |  |  |
| Overall navigation score | 0.30 | 0.96 | 0.33 | 0.96 | 0.23 | 0.96 | 0.03 | 0.96 |
| Scene recognition | 0.01 | 0.98 | 0.00 | 0.98 | 0.05 | 0.96 | 0.19 | 0.96 |
| Proximity judgements | 1.42 | 0.96 | 1.42 | 0.96 | 1.16 | 0.96 | 0.08 | 0.96 |
| Route knowledge | 0.06 | 0.96 | 0.31 | 0.96 | 0.83 | 0.96 | 5.45 | 0.37 |
| Sketch map | 0.11 | 0.96 | 0.10 | 0.96 | 0.10 | 0.96 | 0.00 | 1.00 |

P values are Benjamini-Hochberg false discovery rate corrected at $p < 0.05$ .

**Table S22.** Comparison of hippocampal grey matter <u>R_2_*</u> when dividing the sample into two groups determined by their <u>median performance</u> on each cognitive task, with age, gender, total intracranial volume and MRI scanner included as covariates.

| Performance variable | Whole hippocampus |  | Anterior hippocampus |  | Posterior hippocampus |  | Posterior/Anterior hippocampus ratio |  |
| --- | --- | --- | --- | --- | --- | --- | --- | --- |
|  | F | p | F | p | F | p | F | p |
| **Scene construction** |  |  |  |  |  |  |  |  |
| Experiential index | 1.72 | 0.72 | 1.52 | 0.74 | 1.18 | 0.75 | 0.22 | 0.88 |
| Spatial references | 4.36 | 0.42 | 2.76 | 0.50 | 4.13 | 0.42 | 0.00 | 0.99 |
| Entities present | 3.06 | 0.50 | 1.38 | 0.75 | 3.71 | 0.47 | 0.36 | 0.88 |
| Sensory descriptions | 0.78 | 0.75 | 1.25 | 0.75 | 0.20 | 0.88 | 0.23 | 0.88 |
| Thoughts/emotions/actions | 0.92 | 0.75 | 0.01 | 0.97 | 3.37 | 0.47 | 2.64 | 0.51 |
| Spatial coherence index | 0.02 | 0.96 | 0.12 | 0.89 | 0.01 | 0.97 | 0.09 | 0.92 |
| **Autobiographical interview** |  |  |  |  |  |  |  |  |
| Internal details | 1.07 | 0.95 | 0.17 | 0.95 | 2.08 | 0.95 | 0.38 | 0.95 |
| Internal events | 0.01 | 0.98 | 0.44 | 0.95 | 0.75 | 0.95 | 2.22 | 0.95 |
| Internal time | 0.08 | 0.95 | 0.74 | 0.95 | 0.14 | 0.95 | 1.49 | 0.95 |
| Internal place | 0.91 | 0.95 | 0.55 | 0.95 | 0.91 | 0.95 | 0.08 | 0.95 |
| Internal perceptual | 1.16 | 0.95 | 0.37 | 0.95 | 1.71 | 0.95 | 0.30 | 0.95 |
| Internal thoughts/emotions | 0.07 | 0.95 | 0.07 | 0.95 | 0.04 | 0.96 | 0.05 | 0.95 |
| Vividness ratings | 0.23 | 0.95 | 0.70 | 0.95 | 0.00 | 1.00 | 0.87 | 0.95 |
| **Future thinking** |  |  |  |  |  |  |  |  |
| Experiential index | 0.04 | 0.97 | 0.21 | 0.95 | 0.01 | 0.98 | 0.17 | 0.95 |
| Spatial references | 2.63 | 0.54 | 4.06 | 0.48 | 0.72 | 0.95 | 1.33 | 0.83 |
| Entities present | 2.67 | 0.54 | 4.27 | 0.48 | 0.67 | 0.95 | 1.97 | 0.74 |
| Sensory descriptions | 0.08 | 0.95 | 0.24 | 0.95 | 0.00 | 1.00 | 0.07 | 0.95 |
| Thoughts/emotions/actions | 0.44 | 0.95 | 0.01 | 0.98 | 1.20 | 0.83 | 0.48 | 0.95 |
| Spatial coherence index | 0.57 | 0.95 | 0.75 | 0.95 | 0.22 | 0.95 | 0.10 | 0.95 |
| **Navigation** |  |  |  |  |  |  |  |  |
| Overall navigation score | 1.25 | 0.96 | 0.16 | 0.96 | 2.53 | 0.89 | 1.40 | 0.96 |
| Scene recognition | 0.82 | 0.96 | 0.14 | 0.96 | 1.53 | 0.96 | 0.73 | 0.96 |
| Proximity judgements | 0.08 | 0.96 | 0.01 | 0.98 | 0.16 | 0.96 | 0.02 | 0.98 |
| Route knowledge | 0.11 | 0.96 | 0.01 | 0.98 | 0.44 | 0.96 | 0.71 | 0.96 |
| Sketch map | 1.38 | 0.96 | 0.06 | 0.96 | 3.50 | 0.72 | 3.11 | 0.79 |

P values are Benjamini-Hochberg false discovery rate corrected at $p < 0.05$ .

**Table S23.** Partial correlations between task performance and hippocampal grey matter <u>MT saturation</u> in the <u>low performing</u> participants only (as determined by a median split for each task) with age, gender, total intracranial volume and MRI scanner as covariates.

| Performance variable | Whole hippocampus |  | Anterior hippocampus |  | Posterior hippocampus |  | Posterior/Anterior hippocampus ratio |  |
| --- | --- | --- | --- | --- | --- | --- | --- | --- |
|  | r | p | r | p | r | p | r | p |
| **Scene construction** |  |  |  |  |  |  |  |  |
| Experiential index | -0.17 | 0.98 | -0.15 | 0.98 | -0.16 | 0.98 | 0.008 | 0.99 |
| Spatial references | -0.22 | 0.59 | -0.15 | 0.98 | -0.24 | 0.52 | -0.12 | 0.98 |
| Entities present | -0.010 | 0.99 | -0.009 | 0.99 | -0.010 | 0.99 | -0.019 | 0.99 |
| Sensory descriptions | -0.12 | 0.98 | -0.033 | 0.98 | -0.19 | 0.96 | -0.18 | 0.96 |
| Thoughts/emotions/actions | -0.17 | 0.98 | -0.16 | 0.98 | -0.14 | 0.98 | 0.10 | 0.98 |
| Spatial coherence index | -0.003 | 0.99 | -0.044 | 0.98 | 0.038 | 0.98 | 0.092 | 0.98 |
| **Autobiographical interview** |  |  |  |  |  |  |  |  |
| Internal details | -0.10 | 0.78 | -0.036 | 0.91 | -0.16 | 0.78 | -0.099 | 0.78 |
| Internal events | 0.14 | 0.78 | 0.15 | 0.78 | 0.11 | 0.78 | -0.063 | 0.79 |
| Internal time | 0.035 | 0.90 | 0.060 | 0.79 | 0.005 | 1.00 | -0.088 | 0.78 |
| Internal place | 0.010 | 0.98 | 0.053 | 0.79 | -0.036 | 0.90 | -0.13 | 0.78 |
| Internal perceptual | -0.009 | 0.99 | 0.086 | 0.78 | -0.11 | 0.78 | -0.21 | 0.72 |
| Internal thoughts/emotions | -0.067 | 0.79 | -0.012 | 0.98 | -0.12 | 0.78 | -0.088 | 0.78 |
| Vividness ratings | -0.084 | 0.78 | -0.004 | 1.00 | -0.16 | 0.78 | -0.16 | 0.78 |
| **Future thinking** |  |  |  |  |  |  |  |  |
| Experiential index | -0.095 | 0.79 | -0.036 | 0.93 | -0.14 | 0.78 | -0.099 | 0.79 |
| Spatial references | -0.087 | 0.79 | -0.032 | 0.93 | -0.13 | 0.78 | -0.11 | 0.78 |
| Entities present | -0.029 | 0.93 | 0.020 | 0.93 | -0.070 | 0.79 | -0.104 | 0.78 |
| Sensory descriptions | -0.16 | 0.78 | -0.13 | 0.78 | -0.18 | 0.78 | -0.018 | 0.93 |
| Thoughts/emotions/actions | -0.012 | 0.95 | -0.013 | 0.95 | -0.009 | 0.95 | -0.008 | 0.95 |
| Spatial coherence index | -0.020 | 0.93 | -0.031 | 0.93 | -0.008 | 0.95 | 0.020 | 0.93 |
| **Navigation** |  |  |  |  |  |  |  |  |
| Overall navigation score | 0.022 | 0.94 | 0.13 | 0.60 | -0.083 | 0.71 | -0.27 | 0.46 |
| Scene recognition | 0.13 | 0.60 | 0.13 | 0.60 | 0.11 | 0.60 | -0.054 | 0.74 |
| Proximity judgements | -0.088 | 0.60 | -0.11 | 0.60 | -0.056 | 0.74 | 0.092 | 0.60 |
| Route knowledge | 0.059 | 0.75 | 0.10 | 0.60 | 0.013 | 0.94 | -0.13 | 0.60 |
| Sketch map | -0.071 | 0.74 | 0.016 | 0.94 | -0.15 | 0.60 | -0.18 | 0.60 |

P values are Benjamini-Hochberg false discovery rate corrected at $p < 0.05$ .

**Table S24.** Partial correlations between task performance and hippocampal grey matter <u>PD</u> in the <u>low performing</u> participants only (as determined by a median split for each task) with age, gender, total intracranial volume and MRI scanner as covariates.

| Performance variable | Whole hippocampus |  | Anterior hippocampus |  | Posterior hippocampus |  | Posterior/Anterior hippocampus ratio |  |
| --- | --- | --- | --- | --- | --- | --- | --- | --- |
|  | r | p | r | p | r | p | r | p |
| **Scene construction** |  |  |  |  |  |  |  |  |
| Experiential index | 0.052 | 0.98 | 0.047 | 0.98 | 0.045 | 0.98 | -0.016 | 0.99 |
| Spatial references | 0.002 | 0.99 | -0.024 | 0.99 | 0.033 | 0.98 | 0.052 | 0.98 |
| Entities present | 0.072 | 0.98 | 0.079 | 0.98 | 0.044 | 0.98 | -0.054 | 0.98 |
| Sensory descriptions | 0.044 | 0.98 | 0.055 | 0.98 | 0.021 | 0.99 | -0.052 | 0.98 |
| Thoughts/emotions/actions | -0.063 | 0.98 | -0.068 | 0.98 | -0.036 | 0.98 | 0.046 | 0.98 |
| Spatial coherence index | 0.069 | 0.98 | 0.11 | 0.98 | 0.002 | 0.99 | -0.13 | 0.98 |
| **Autobiographical interview** |  |  |  |  |  |  |  |  |
| Internal details | -0.076 | 0.79 | -0.094 | 0.78 | -0.029 | 0.94 | 0.086 | 0.78 |
| Internal events | -0.092 | 0.78 | -0.057 | 0.79 | -0.11 | 0.78 | -0.013 | 0.98 |
| Internal time | -0.059 | 0.79 | -0.088 | 0.78 | 0.00 | 1.00 | 0.092 | 0.78 |
| Internal place | -0.12 | 0.78 | -0.10 | 0.78 | -0.13 | 0.78 | 0.033 | 0.91 |
| Internal perceptual | -0.13 | 0.78 | -0.13 | 0.78 | -0.086 | 0.78 | 0.089 | 0.78 |
| Internal thoughts/emotions | 0.12 | 0.78 | 0.089 | 0.78 | 0.12 | 0.78 | -0.006 | 1.00 |
| Vividness ratings | -0.20 | 0.77 | -0.15 | 0.78 | -0.22 | 0.72 | -0.002 | 1.00 |
| **Future thinking** |  |  |  |  |  |  |  |  |
| Experiential index | 0.073 | 0.79 | 0.12 | 0.78 | 0.00 | 1.00 | -0.15 | 0.78 |
| Spatial references | 0.11 | 0.78 | 0.098 | 0.78 | 0.098 | 0.78 | -0.040 | 0.91 |
| Entities present | 0.041 | 0.91 | 0.041 | 0.91 | 0.030 | 0.93 | -0.026 | 0.93 |
| Sensory descriptions | 0.10 | 0.78 | 0.10 | 0.78 | 0.071 | 0.79 | -0.064 | 0.82 |
| Thoughts/emotions/actions | 0.085 | 0.79 | 0.12 | 0.78 | 0.008 | 0.95 | -0.13 | 0.78 |
| Spatial coherence index | 0.083 | 0.79 | 0.12 | 0.78 | 0.022 | 0.93 | -0.12 | 0.78 |
| **Navigation** |  |  |  |  |  |  |  |  |
| Overall navigation score | -0.066 | 0.74 | -0.11 | 0.60 | 0.002 | 0.99 | 0.12 | 0.60 |
| Scene recognition | -0.15 | 0.60 | -0.13 | 0.60 | -0.12 | 0.60 | 0.055 | 0.74 |
| Proximity judgements | 0.085 | 0.60 | 0.023 | 0.93 | 0.13 | 0.60 | 0.085 | 0.60 |
| Route knowledge | -0.12 | 0.60 | -0.098 | 0.60 | -0.11 | 0.60 | 0.017 | 0.94 |
| Sketch map | 0.003 | 0.98 | -0.053 | 0.78 | 0.062 | 0.74 | 0.11 | 0.60 |

P values are Benjamini-Hochberg false discovery rate corrected at $p < 0.05$ .

**Table S25.**
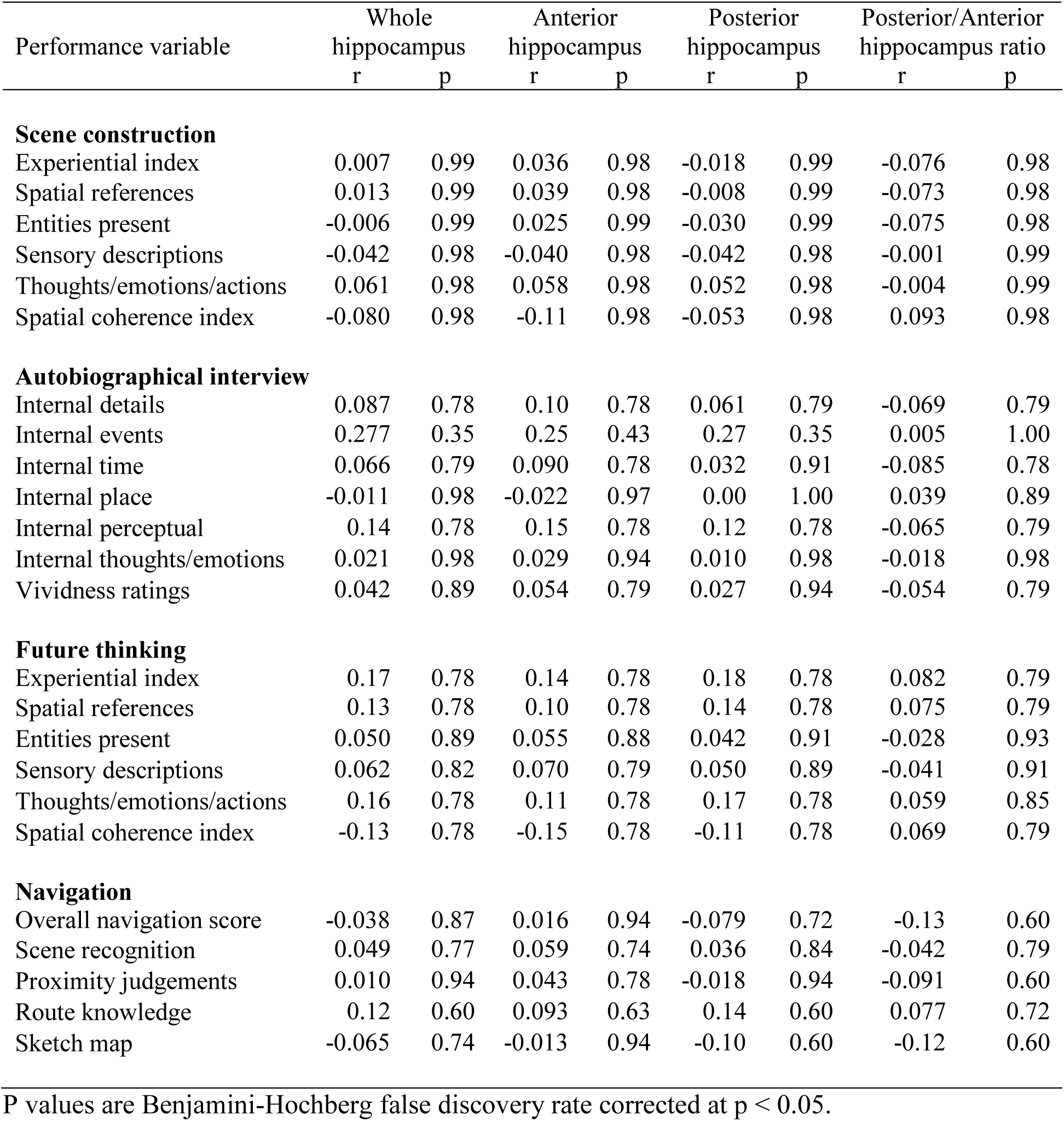
Partial correlations between task performance and hippocampal grey matter <u>R_1_</u> in the <u>low performing</u> participants only (as determined by a median split for each task) with age, gender, total intracranial volume and MRI scanner as covariates.

| Performance variable | Whole hippocampus |  | Anterior hippocampus |  | Posterior hippocampus |  | Posterior/Anterior hippocampus ratio |  |
| --- | --- | --- | --- | --- | --- | --- | --- | --- |
|  | r | p | r | p | r | p | r | p |
| **Scene construction** |  |  |  |  |  |  |  |  |
| Experiential index | 0.007 | 0.99 | 0.036 | 0.98 | -0.018 | 0.99 | -0.076 | 0.98 |
| Spatial references | 0.013 | 0.99 | 0.039 | 0.98 | -0.008 | 0.99 | -0.073 | 0.98 |
| Entities present | -0.006 | 0.99 | 0.025 | 0.99 | -0.030 | 0.99 | -0.075 | 0.98 |
| Sensory descriptions | -0.042 | 0.98 | -0.040 | 0.98 | -0.042 | 0.98 | -0.001 | 0.99 |
| Thoughts/emotions/actions | 0.061 | 0.98 | 0.058 | 0.98 | 0.052 | 0.98 | -0.004 | 0.99 |
| Spatial coherence index | -0.080 | 0.98 | -0.11 | 0.98 | -0.053 | 0.98 | 0.093 | 0.98 |
| **Autobiographical interview** |  |  |  |  |  |  |  |  |
| Internal details | 0.087 | 0.78 | 0.10 | 0.78 | 0.061 | 0.79 | -0.069 | 0.79 |
| Internal events | 0.277 | 0.35 | 0.25 | 0.43 | 0.27 | 0.35 | 0.005 | 1.00 |
| Internal time | 0.066 | 0.79 | 0.090 | 0.78 | 0.032 | 0.91 | -0.085 | 0.78 |
| Internal place | -0.011 | 0.98 | -0.022 | 0.97 | 0.00 | 1.00 | 0.039 | 0.89 |
| Internal perceptual | 0.14 | 0.78 | 0.15 | 0.78 | 0.12 | 0.78 | -0.065 | 0.79 |
| Internal thoughts/emotions | 0.021 | 0.98 | 0.029 | 0.94 | 0.010 | 0.98 | -0.018 | 0.98 |
| Vividness ratings | 0.042 | 0.89 | 0.054 | 0.79 | 0.027 | 0.94 | -0.054 | 0.79 |
| **Future thinking** |  |  |  |  |  |  |  |  |
| Experiential index | 0.17 | 0.78 | 0.14 | 0.78 | 0.18 | 0.78 | 0.082 | 0.79 |
| Spatial references | 0.13 | 0.78 | 0.10 | 0.78 | 0.14 | 0.78 | 0.075 | 0.79 |
| Entities present | 0.050 | 0.89 | 0.055 | 0.88 | 0.042 | 0.91 | -0.028 | 0.93 |
| Sensory descriptions | 0.062 | 0.82 | 0.070 | 0.79 | 0.050 | 0.89 | -0.041 | 0.91 |
| Thoughts/emotions/actions | 0.16 | 0.78 | 0.11 | 0.78 | 0.17 | 0.78 | 0.059 | 0.85 |
| Spatial coherence index | -0.13 | 0.78 | -0.15 | 0.78 | -0.11 | 0.78 | 0.069 | 0.79 |
| **Navigation** |  |  |  |  |  |  |  |  |
| Overall navigation score | -0.038 | 0.87 | 0.016 | 0.94 | -0.079 | 0.72 | -0.13 | 0.60 |
| Scene recognition | 0.049 | 0.77 | 0.059 | 0.74 | 0.036 | 0.84 | -0.042 | 0.79 |
| Proximity judgements | 0.010 | 0.94 | 0.043 | 0.78 | -0.018 | 0.94 | -0.091 | 0.60 |
| Route knowledge | 0.12 | 0.60 | 0.093 | 0.63 | 0.14 | 0.60 | 0.077 | 0.72 |
| Sketch map | -0.065 | 0.74 | -0.013 | 0.94 | -0.10 | 0.60 | -0.12 | 0.60 |

P values are Benjamini-Hochberg false discovery rate corrected at $p < 0.05$ .

**Table S26.**
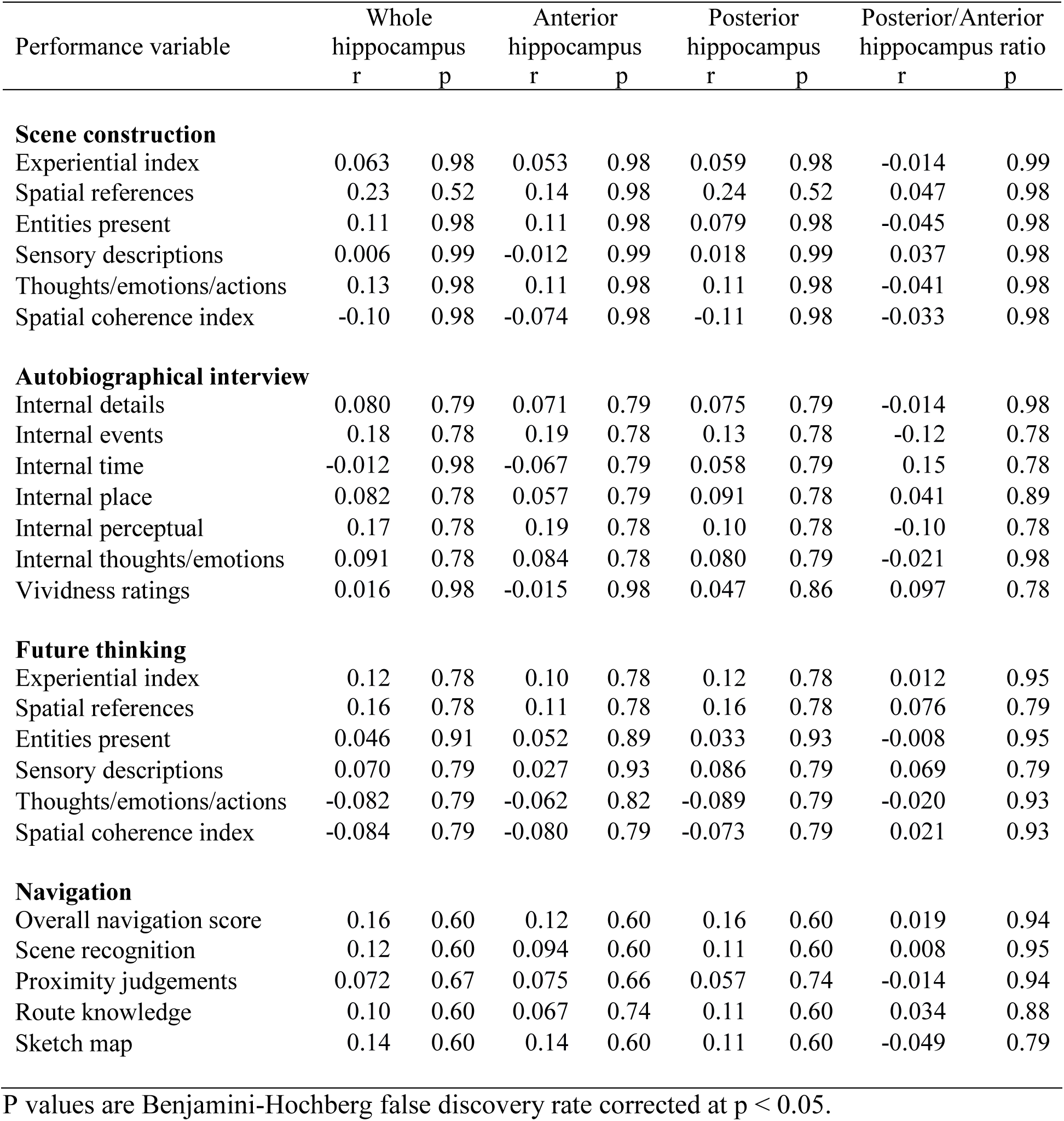
Partial correlations between task performance and hippocampal grey matter <u>R_2_*</u> in the low performing participants only (as determined by a median split for each task) with age, gender, total intracranial volume and MRI scanner as covariates.

**Table S27.** Partial correlations between performance and hippocampal grey matter <u>MT saturation</u> in the <u>high performing</u> participants only (as determined by a median split for each task) with age, gender, total intracranial volume and MRI scanner as covariates.

| Performance variable | Whole hippocampus |  | Anterior hippocampus |  | Posterior hippocampus |  | Posterior/Anterior hippocampus ratio |  |
| --- | --- | --- | --- | --- | --- | --- | --- | --- |
|  | r | p | r | p | r | p | r | p |
| **Scene construction** |  |  |  |  |  |  |  |  |
| Experiential index | -0.13 | 0.77 | -0.14 | 0.77 | -0.10 | 0.77 | 0.090 | 0.77 |
| Spatial references | 0.001 | 1.00 | -0.045 | 0.91 | 0.051 | 0.91 | 0.11 | 0.77 |
| Entities present | -0.12 | 0.77 | -0.13 | 0.77 | -0.099 | 0.77 | 0.099 | 0.77 |
| Sensory descriptions | -0.036 | 0.96 | -0.095 | 0.77 | 0.027 | 0.99 | 0.17 | 0.77 |
| Thoughts/emotions/actions | -0.18 | 0.77 | -0.22 | 0.77 | -0.12 | 0.77 | 0.19 | 0.77 |
| Spatial coherence index | -0.13 | 0.77 | -0.14 | 0.77 | -0.10 | 0.77 | 0.11 | 0.77 |
| **Autobiographical interview** |  |  |  |  |  |  |  |  |
| Internal details | 0.009 | 0.99 | -0.013 | 0.99 | 0.027 | 0.95 | 0.047 | 0.93 |
| Internal events | 0.086 | 0.81 | 0.088 | 0.81 | 0.074 | 0.81 | -0.043 | 0.93 |
| Internal time | 0.14 | 0.75 | 0.16 | 0.75 | 0.10 | 0.81 | -0.091 | 0.81 |
| Internal place | -0.073 | 0.81 | -0.10 | 0.81 | -0.040 | 0.94 | 0.078 | 0.81 |
| Internal perceptual | 0.016 | 0.99 | 0.011 | 0.99 | 0.019 | 0.99 | 0.008 | 0.99 |
| Internal thoughts/emotions | -0.032 | 0.94 | -0.062 | 0.82 | -0.001 | 1.00 | 0.096 | 0.81 |
| Vividness ratings | -0.043 | 0.93 | -0.076 | 0.81 | -0.011 | 0.99 | 0.087 | 0.81 |
| **Future thinking** |  |  |  |  |  |  |  |  |
| Experiential index | -0.28 | 0.17 | -0.32 | 0.11 | -0.20 | 0.36 | 0.25 | 0.28 |
| Spatial references | -0.17 | 0.49 | -0.20 | 0.39 | -0.122 | 0.74 | 0.15 | 0.59 |
| Entities present | -0.22 | 0.34 | -0.23 | 0.32 | -0.19 | 0.39 | 0.15 | 0.54 |
| Sensory descriptions | -0.10 | 0.79 | -0.089 | 0.80 | -0.097 | 0.79 | 0.061 | 0.84 |
| Thoughts/emotions/actions | 0.21 | 0.35 | 0.16 | 0.49 | 0.24 | 0.29 | 0.053 | 0.84 |
| Spatial coherence index | -0.12 | 0.74 | -0.098 | 0.79 | -0.12 | 0.74 | 0.031 | 0.88 |
| **Navigation** |  |  |  |  |  |  |  |  |
| Overall navigation score | 0.077 | 0.99 | 0.059 | 0.99 | 0.084 | 0.99 | 0.000 | 1.00 |
| Scene recognition | 0.22 | 0.99 | 0.24 | 0.99 | 0.17 | 0.99 | -0.14 | 0.99 |
| Proximity judgements | -0.090 | 0.99 | -0.055 | 0.99 | -0.12 | 0.99 | -0.047 | 0.99 |
| Route knowledge | -0.10 | 0.99 | -0.11 | 0.99 | -0.069 | 0.99 | 0.065 | 0.99 |
| Sketch map | 0.062 | 0.99 | 0.017 | 1.00 | 0.097 | 0.99 | 0.074 | 0.99 |

P values are Benjamini-Hochberg false discovery rate corrected at $p < 0.05$ .

**Table S28.**
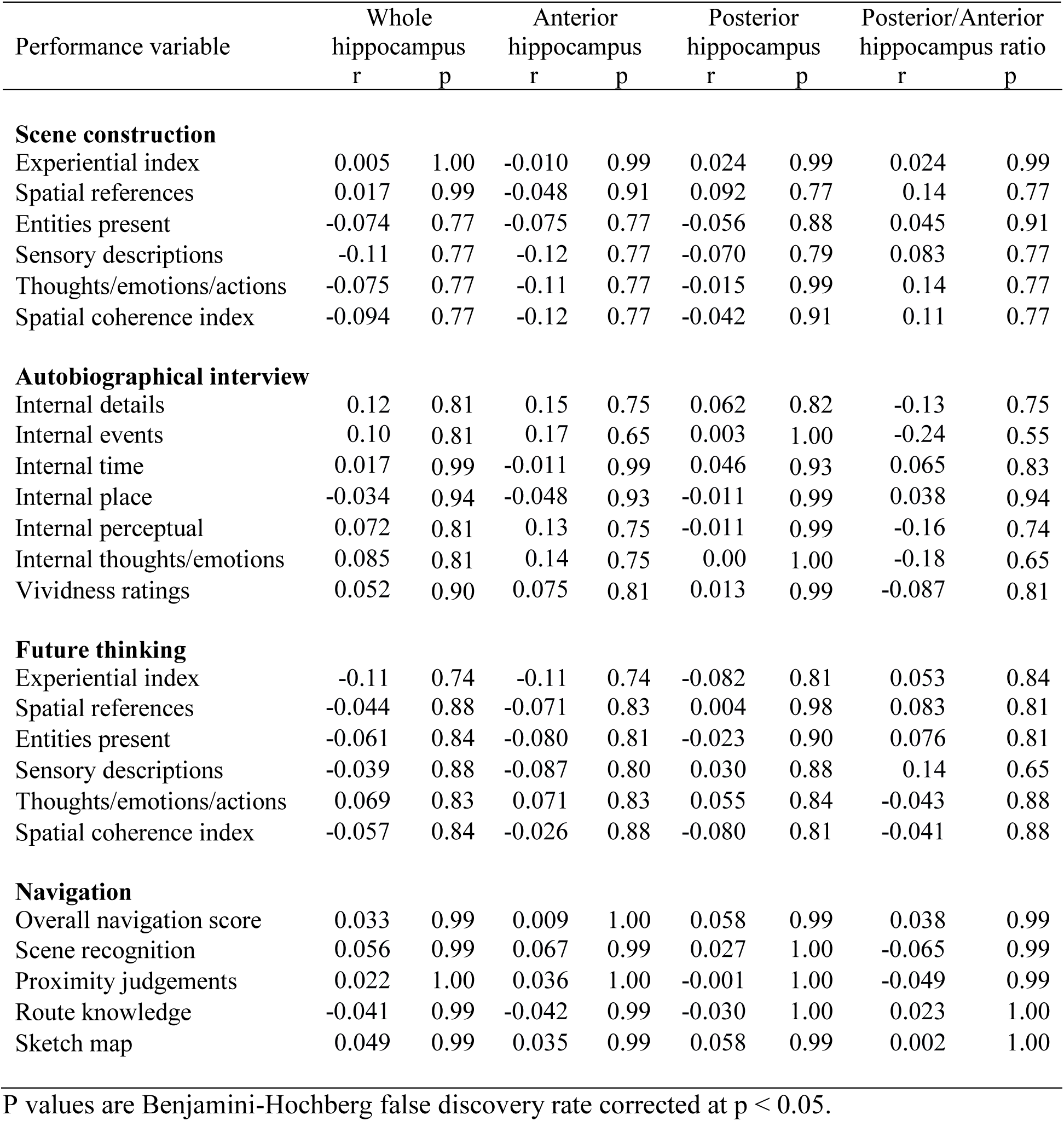
Partial correlations between performance and hippocampal grey matter <u>PD</u> in the <u>high performing</u> participants only (as determined by a median split for each task) with age, gender, total intracranial volume and MRI scanner as covariates.

| Performance variable | Whole hippocampus |  | Anterior hippocampus |  | Posterior hippocampus |  | Posterior/Anterior hippocampus ratio |  |
| --- | --- | --- | --- | --- | --- | --- | --- | --- |
|  | r | p | r | p | r | p | r | p |
| **Scene construction** |  |  |  |  |  |  |  |  |
| Experiential index | 0.005 | 1.00 | -0.010 | 0.99 | 0.024 | 0.99 | 0.024 | 0.99 |
| Spatial references | 0.017 | 0.99 | -0.048 | 0.91 | 0.092 | 0.77 | 0.14 | 0.77 |
| Entities present | -0.074 | 0.77 | -0.075 | 0.77 | -0.056 | 0.88 | 0.045 | 0.91 |
| Sensory descriptions | -0.11 | 0.77 | -0.12 | 0.77 | -0.070 | 0.79 | 0.083 | 0.77 |
| Thoughts/emotions/actions | -0.075 | 0.77 | -0.11 | 0.77 | -0.015 | 0.99 | 0.14 | 0.77 |
| Spatial coherence index | -0.094 | 0.77 | -0.12 | 0.77 | -0.042 | 0.91 | 0.11 | 0.77 |
| **Autobiographical interview** |  |  |  |  |  |  |  |  |
| Internal details | 0.12 | 0.81 | 0.15 | 0.75 | 0.062 | 0.82 | -0.13 | 0.75 |
| Internal events | 0.10 | 0.81 | 0.17 | 0.65 | 0.003 | 1.00 | -0.24 | 0.55 |
| Internal time | 0.017 | 0.99 | -0.011 | 0.99 | 0.046 | 0.93 | 0.065 | 0.83 |
| Internal place | -0.034 | 0.94 | -0.048 | 0.93 | -0.011 | 0.99 | 0.038 | 0.94 |
| Internal perceptual | 0.072 | 0.81 | 0.13 | 0.75 | -0.011 | 0.99 | -0.16 | 0.74 |
| Internal thoughts/emotions | 0.085 | 0.81 | 0.14 | 0.75 | 0.00 | 1.00 | -0.18 | 0.65 |
| Vividness ratings | 0.052 | 0.90 | 0.075 | 0.81 | 0.013 | 0.99 | -0.087 | 0.81 |
| **Future thinking** |  |  |  |  |  |  |  |  |
| Experiential index | -0.11 | 0.74 | -0.11 | 0.74 | -0.082 | 0.81 | 0.053 | 0.84 |
| Spatial references | -0.044 | 0.88 | -0.071 | 0.83 | 0.004 | 0.98 | 0.083 | 0.81 |
| Entities present | -0.061 | 0.84 | -0.080 | 0.81 | -0.023 | 0.90 | 0.076 | 0.81 |
| Sensory descriptions | -0.039 | 0.88 | -0.087 | 0.80 | 0.030 | 0.88 | 0.14 | 0.65 |
| Thoughts/emotions/actions | 0.069 | 0.83 | 0.071 | 0.83 | 0.055 | 0.84 | -0.043 | 0.88 |
| Spatial coherence index | -0.057 | 0.84 | -0.026 | 0.88 | -0.080 | 0.81 | -0.041 | 0.88 |
| **Navigation** |  |  |  |  |  |  |  |  |
| Overall navigation score | 0.033 | 0.99 | 0.009 | 1.00 | 0.058 | 0.99 | 0.038 | 0.99 |
| Scene recognition | 0.056 | 0.99 | 0.067 | 0.99 | 0.027 | 1.00 | -0.065 | 0.99 |
| Proximity judgements | 0.022 | 1.00 | 0.036 | 1.00 | -0.001 | 1.00 | -0.049 | 0.99 |
| Route knowledge | -0.041 | 0.99 | -0.042 | 0.99 | -0.030 | 1.00 | 0.023 | 1.00 |
| Sketch map | 0.049 | 0.99 | 0.035 | 0.99 | 0.058 | 0.99 | 0.002 | 1.00 |

P values are Benjamini-Hochberg false discovery rate corrected at $p < 0.05$ .

**Table S29.** Partial correlations between performance and hippocampal grey matter <u>R_1_</u> in the <u>high performing</u> participants only (as determined by a median split for each task) with age, gender, total intracranial volume and MRI scanner as covariates.

| Performance variable | Whole hippocampus |  | Anterior hippocampus |  | Posterior hippocampus |  | Posterior/Anterior hippocampus ratio |  |
| --- | --- | --- | --- | --- | --- | --- | --- | --- |
|  | r | p | r | p | r | p | r | p |
| **Scene construction** |  |  |  |  |  |  |  |  |
| Experiential index | -0.008 | 0.99 | -0.004 | 1.00 | -0.010 | 0.99 | -0.019 | 0.99 |
| Spatial references | -0.007 | 0.99 | 0.000 | 1.00 | -0.013 | 0.99 | -0.019 | 0.99 |
| Entities present | -0.055 | 0.88 | -0.020 | 0.99 | -0.080 | 0.77 | -0.10 | 0.77 |
| Sensory descriptions | 0.17 | 0.77 | 0.15 | 0.77 | 0.17 | 0.77 | 0.008 | 0.99 |
| Thoughts/emotions/actions | 0.13 | 0.77 | 0.15 | 0.77 | 0.11 | 0.77 | -0.095 | 0.77 |
| Spatial coherence index | 0.075 | 0.77 | 0.083 | 0.77 | 0.058 | 0.88 | -0.048 | 0.91 |
| **Autobiographical interview** |  |  |  |  |  |  |  |  |
| Internal details | -0.084 | 0.81 | -0.082 | 0.81 | -0.080 | 0.81 | 0.006 | 1.00 |
| Internal events | -0.056 | 0.86 | -0.081 | 0.81 | -0.031 | 0.94 | 0.098 | 0.81 |
| Internal time | -0.12 | 0.81 | -0.073 | 0.82 | -0.15 | 0.75 | -0.16 | 0.75 |
| Internal place | 0.15 | 0.75 | 0.11 | 0.81 | 0.16 | 0.75 | 0.077 | 0.81 |
| Internal perceptual | 0.090 | 0.81 | 0.068 | 0.81 | 0.097 | 0.81 | 0.038 | 0.94 |
| Internal thoughts/emotions | -0.029 | 0.94 | -0.064 | 0.82 | 0.001 | 1.00 | 0.13 | 0.75 |
| Vividness ratings | -0.21 | 0.65 | -0.18 | 0.65 | -0.20 | 0.65 | -0.072 | 0.81 |
| **Future thinking** |  |  |  |  |  |  |  |  |
| Experiential index | 0.042 | 0.88 | 0.047 | 0.88 | 0.033 | 0.88 | -0.026 | 0.88 |
| Spatial references | 0.12 | 0.74 | 0.12 | 0.74 | 0.094 | 0.80 | -0.037 | 0.88 |
| Entities present | 0.066 | 0.84 | 0.096 | 0.79 | 0.032 | 0.88 | -0.088 | 0.80 |
| Sensory descriptions | 0.17 | 0.49 | 0.18 | 0.47 | 0.14 | 0.61 | -0.057 | 0.84 |
| Thoughts/emotions/actions | -0.065 | 0.84 | -0.072 | 0.83 | -0.055 | 0.84 | 0.034 | 0.88 |
| Spatial coherence index | 0.000 | 1.00 | -0.005 | 0.98 | 0.004 | 0.98 | 0.014 | 0.95 |
| **Navigation** |  |  |  |  |  |  |  |  |
| Overall navigation score | -0.11 | 0.99 | -0.10 | 0.99 | -0.11 | 0.99 | -0.013 | 1.00 |
| Scene recognition | -0.13 | 0.99 | -0.074 | 0.99 | -0.15 | 0.99 | -0.11 | 0.99 |
| Proximity judgements | -0.15 | 0.99 | -0.13 | 0.99 | -0.15 | 0.99 | 0.001 | 1.00 |
| Route knowledge | -0.054 | 0.99 | -0.064 | 0.99 | -0.039 | 0.99 | 0.023 | 1.00 |
| Sketch map | -0.059 | 0.99 | -0.058 | 0.99 | -0.056 | 0.99 | 0.000 | 1.00 |

P values are Benjamini-Hochberg false discovery rate corrected at $p < 0.05$ .

**Table S30.** Partial correlations between performance and hippocampal grey matter <u>R_2_*</u> in the <u>high performing</u> participants only (as determined by a median split for each task) with age, gender, total intracranial volume and MRI scanner as covariates.

| Performance variable | Whole hippocampus |  | Anterior hippocampus |  | Posterior hippocampus |  | Posterior/Anterior hippocampus ratio |  |
| --- | --- | --- | --- | --- | --- | --- | --- | --- |
|  | r | p | r | p | r | p | r | p |
| **Scene construction** |  |  |  |  |  |  |  |  |
| Experiential index | -0.057 | 0.88 | -0.11 | 0.77 | 0.008 | 0.99 | 0.13 | 0.77 |
| Spatial references | -0.080 | 0.77 | -0.083 | 0.77 | -0.062 | 0.86 | 0.021 | 0.99 |
| Entities present | -0.018 | 0.99 | -0.074 | 0.77 | 0.044 | 0.91 | 0.15 | 0.77 |
| Sensory descriptions | -0.002 | 1.00 | -0.077 | 0.77 | 0.091 | 0.77 | 0.15 | 0.77 |
| Thoughts/emotions/actions | 0.009 | 0.99 | -0.085 | 0.77 | 0.081 | 0.77 | 0.18 | 0.77 |
| Spatial coherence index | 0.023 | 0.99 | 0.13 | 0.77 | -0.12 | 0.77 | -0.22 | 0.77 |
| **Autobiographical interview** |  |  |  |  |  |  |  |  |
| Internal details | -0.16 | 0.75 | -0.15 | 0.75 | -0.13 | 0.75 | 0.010 | 0.99 |
| Internal events | -0.081 | 0.81 | -0.072 | 0.81 | -0.069 | 0.81 | -0.001 | 1.00 |
| Internal time | -0.10 | 0.81 | -0.014 | 0.99 | -0.15 | 0.75 | -0.15 | 0.75 |
| Internal place | -0.038 | 0.94 | -0.065 | 0.82 | -0.002 | 1.00 | 0.070 | 0.82 |
| Internal perceptual | -0.069 | 0.81 | -0.031 | 0.94 | -0.094 | 0.81 | -0.084 | 0.81 |
| Internal thoughts/emotions | -0.10 | 0.81 | -0.15 | 0.75 | -0.044 | 0.93 | 0.11 | 0.81 |
| Vividness ratings | -0.14 | 0.75 | 0.029 | 0.94 | -0.26 | 0.55 | -0.28 | 0.55 |
| **Future thinking** |  |  |  |  |  |  |  |  |
| Experiential index | 0.18 | 0.39 | 0.16 | 0.49 | 0.16 | 0.49 | -0.099 | 0.79 |
| Spatial references | 0.091 | 0.80 | 0.021 | 0.91 | 0.16 | 0.54 | 0.065 | 0.84 |
| Entities present | 0.077 | 0.81 | 0.044 | 0.88 | 0.10 | 0.79 | 0.038 | 0.88 |
| Sensory descriptions | 0.20 | 0.39 | 0.19 | 0.39 | 0.17 | 0.49 | -0.087 | 0.80 |
| Thoughts/emotions/actions | 0.007 | 0.98 | 0.027 | 0.88 | -0.010 | 0.98 | -0.033 | 0.88 |
| Spatial coherence index | 0.25 | 0.29 | 0.28 | 0.17 | 0.13 | 0.66 | -0.22 | 0.35 |
| **Navigation** |  |  |  |  |  |  |  |  |
| Overall navigation score | -0.085 | 0.99 | -0.092 | 0.99 | -0.063 | 0.99 | 0.060 | 0.99 |
| Scene recognition | -0.023 | 1.00 | -0.035 | 1.00 | -0.008 | 1.00 | 0.024 | 1.00 |
| Proximity judgements | -0.097 | 0.99 | 0.003 | 1.00 | -0.21 | 0.99 | -0.15 | 0.99 |
| Route knowledge | -0.11 | 0.99 | -0.15 | 0.99 | -0.046 | 0.99 | 0.134 | 0.99 |
| Sketch map | -0.084 | 0.99 | -0.068 | 0.99 | -0.084 | 0.99 | 0.00 | 1.00 |

P values are Benjamini-Hochberg false discovery rate corrected at $p < 0.05$ .

**Table S31.**
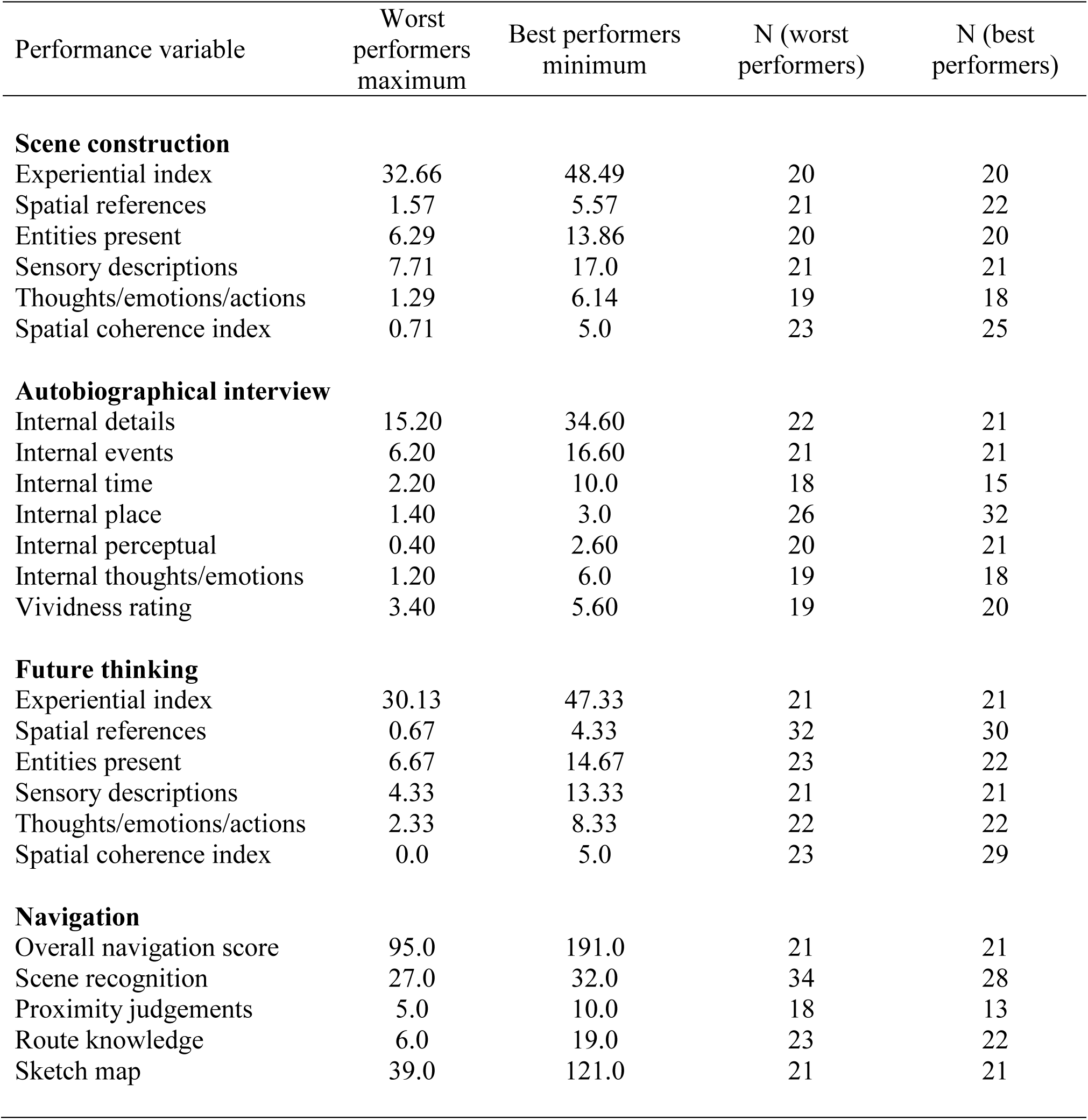
Details of the groups created for each task when taking only the best and worst performers.

**Table S32.**
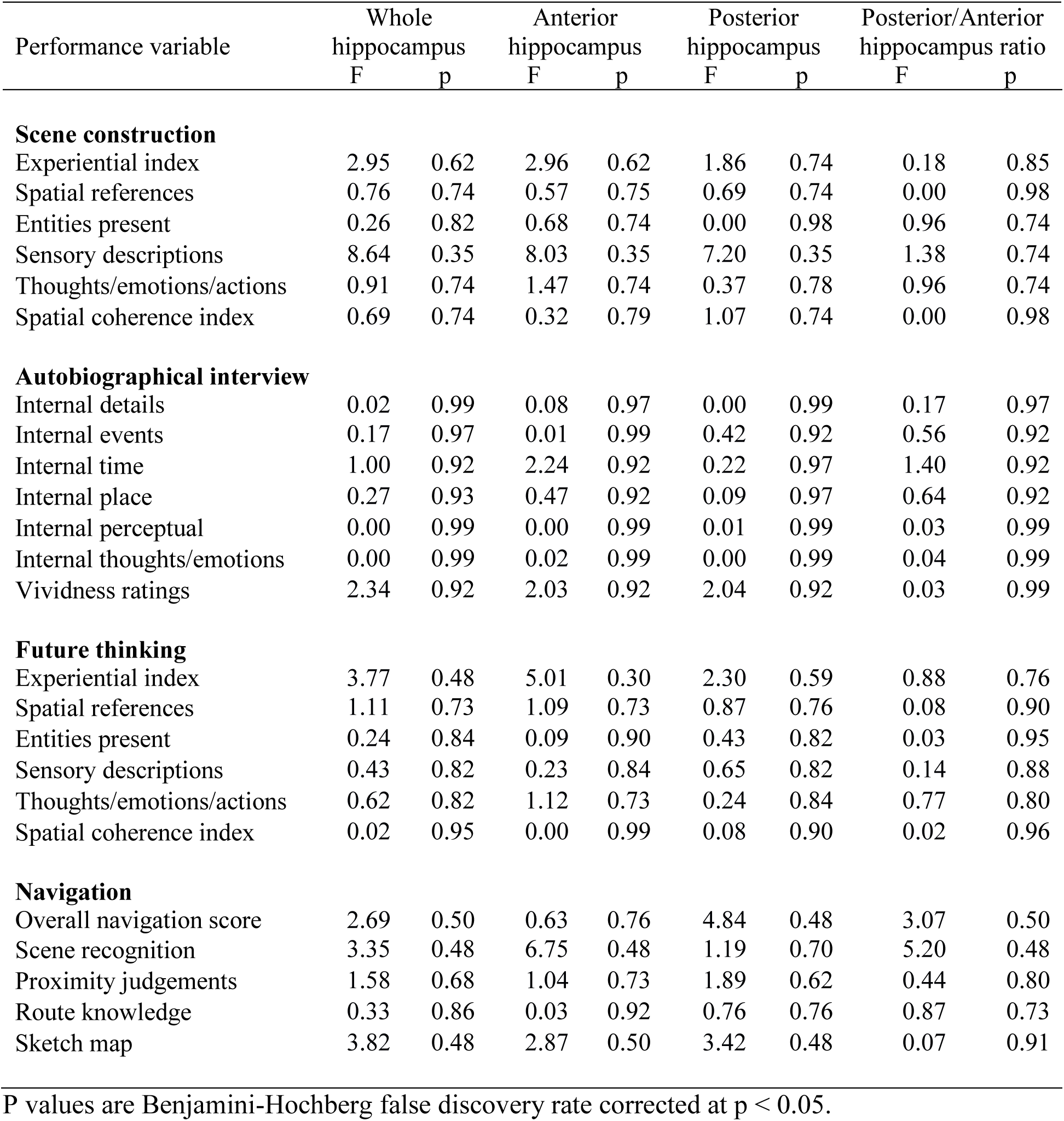
Comparison of hippocampal grey matter <u>MT saturation</u> when taking the <u>best and worst</u> performing participants for task, with age, gender, total intracranial volume and MRI scanner included as covariates.

| Performance variable | Whole hippocampus |  | Anterior hippocampus |  | Posterior hippocampus |  | Posterior/Anterior hippocampus ratio |  |
| --- | --- | --- | --- | --- | --- | --- | --- | --- |
|  | F | p | F | p | F | p | F | p |
| **Scene construction** |  |  |  |  |  |  |  |  |
| Experiential index | 2.95 | 0.62 | 2.96 | 0.62 | 1.86 | 0.74 | 0.18 | 0.85 |
| Spatial references | 0.76 | 0.74 | 0.57 | 0.75 | 0.69 | 0.74 | 0.00 | 0.98 |
| Entities present | 0.26 | 0.82 | 0.68 | 0.74 | 0.00 | 0.98 | 0.96 | 0.74 |
| Sensory descriptions | 8.64 | 0.35 | 8.03 | 0.35 | 7.20 | 0.35 | 1.38 | 0.74 |
| Thoughts/emotions/actions | 0.91 | 0.74 | 1.47 | 0.74 | 0.37 | 0.78 | 0.96 | 0.74 |
| Spatial coherence index | 0.69 | 0.74 | 0.32 | 0.79 | 1.07 | 0.74 | 0.00 | 0.98 |
| **Autobiographical interview** |  |  |  |  |  |  |  |  |
| Internal details | 0.02 | 0.99 | 0.08 | 0.97 | 0.00 | 0.99 | 0.17 | 0.97 |
| Internal events | 0.17 | 0.97 | 0.01 | 0.99 | 0.42 | 0.92 | 0.56 | 0.92 |
| Internal time | 1.00 | 0.92 | 2.24 | 0.92 | 0.22 | 0.97 | 1.40 | 0.92 |
| Internal place | 0.27 | 0.93 | 0.47 | 0.92 | 0.09 | 0.97 | 0.64 | 0.92 |
| Internal perceptual | 0.00 | 0.99 | 0.00 | 0.99 | 0.01 | 0.99 | 0.03 | 0.99 |
| Internal thoughts/emotions | 0.00 | 0.99 | 0.02 | 0.99 | 0.00 | 0.99 | 0.04 | 0.99 |
| Vividness ratings | 2.34 | 0.92 | 2.03 | 0.92 | 2.04 | 0.92 | 0.03 | 0.99 |
| **Future thinking** |  |  |  |  |  |  |  |  |
| Experiential index | 3.77 | 0.48 | 5.01 | 0.30 | 2.30 | 0.59 | 0.88 | 0.76 |
| Spatial references | 1.11 | 0.73 | 1.09 | 0.73 | 0.87 | 0.76 | 0.08 | 0.90 |
| Entities present | 0.24 | 0.84 | 0.09 | 0.90 | 0.43 | 0.82 | 0.03 | 0.95 |
| Sensory descriptions | 0.43 | 0.82 | 0.23 | 0.84 | 0.65 | 0.82 | 0.14 | 0.88 |
| Thoughts/emotions/actions | 0.62 | 0.82 | 1.12 | 0.73 | 0.24 | 0.84 | 0.77 | 0.80 |
| Spatial coherence index | 0.02 | 0.95 | 0.00 | 0.99 | 0.08 | 0.90 | 0.02 | 0.96 |
| **Navigation** |  |  |  |  |  |  |  |  |
| Overall navigation score | 2.69 | 0.50 | 0.63 | 0.76 | 4.84 | 0.48 | 3.07 | 0.50 |
| Scene recognition | 3.35 | 0.48 | 6.75 | 0.48 | 1.19 | 0.70 | 5.20 | 0.48 |
| Proximity judgements | 1.58 | 0.68 | 1.04 | 0.73 | 1.89 | 0.62 | 0.44 | 0.80 |
| Route knowledge | 0.33 | 0.86 | 0.03 | 0.92 | 0.76 | 0.76 | 0.87 | 0.73 |
| Sketch map | 3.82 | 0.48 | 2.87 | 0.50 | 3.42 | 0.48 | 0.07 | 0.91 |

P values are Benjamini-Hochberg false discovery rate corrected at $p < 0.05$ .

**Table S33.**
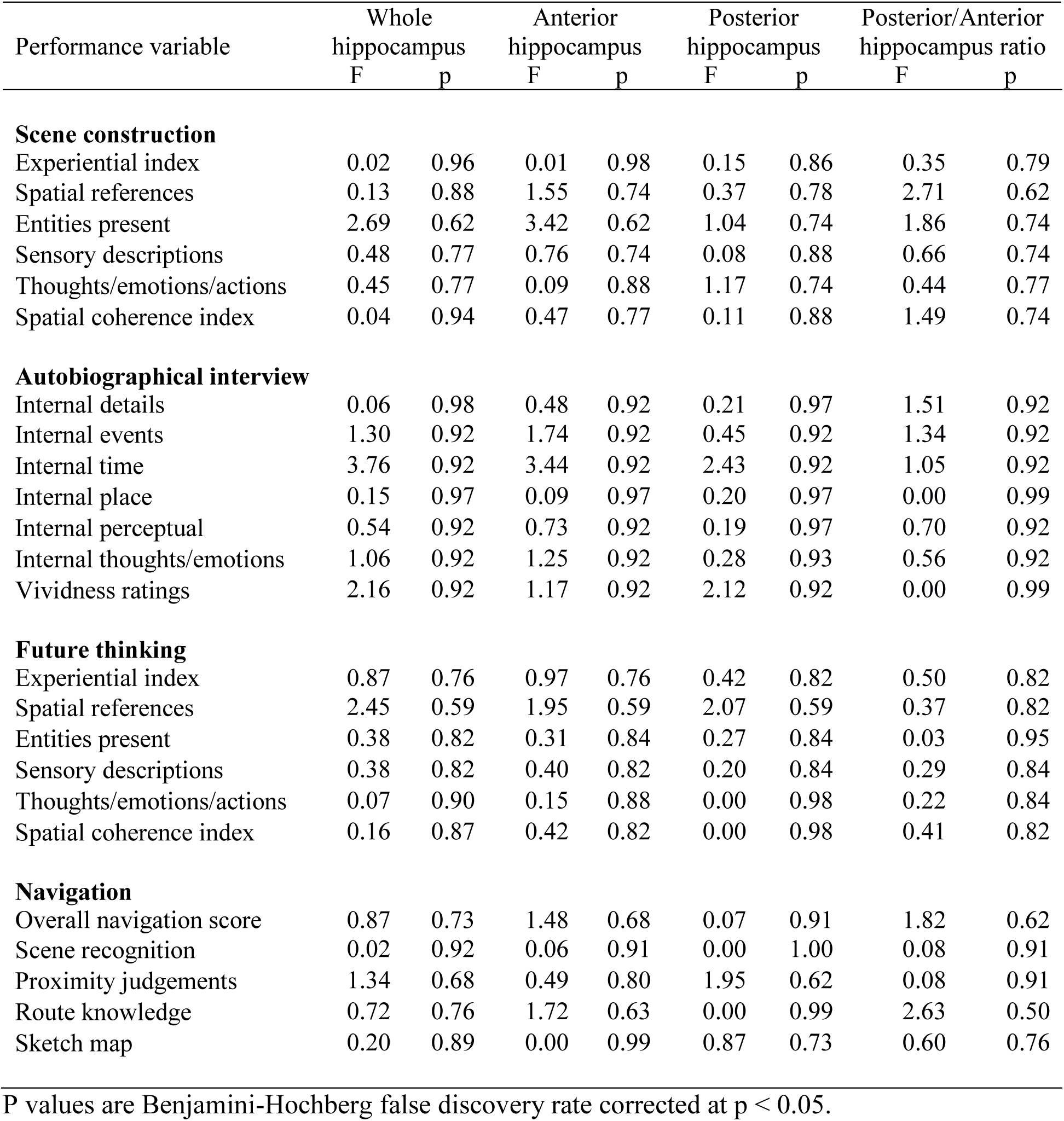
Comparison of hippocampal grey matter <u>PD</u> when taking the <u>best and worst</u> performing participants for task, with age, gender, total intracranial volume and MRI scanner included as covariates.

| Performance variable | Whole hippocampus |  | Anterior hippocampus |  | Posterior hippocampus |  | Posterior/Anterior hippocampus ratio |  |
| --- | --- | --- | --- | --- | --- | --- | --- | --- |
|  | F | p | F | p | F | p | F | p |
| **Scene construction** |  |  |  |  |  |  |  |  |
| Experiential index | 0.02 | 0.96 | 0.01 | 0.98 | 0.15 | 0.86 | 0.35 | 0.79 |
| Spatial references | 0.13 | 0.88 | 1.55 | 0.74 | 0.37 | 0.78 | 2.71 | 0.62 |
| Entities present | 2.69 | 0.62 | 3.42 | 0.62 | 1.04 | 0.74 | 1.86 | 0.74 |
| Sensory descriptions | 0.48 | 0.77 | 0.76 | 0.74 | 0.08 | 0.88 | 0.66 | 0.74 |
| Thoughts/emotions/actions | 0.45 | 0.77 | 0.09 | 0.88 | 1.17 | 0.74 | 0.44 | 0.77 |
| Spatial coherence index | 0.04 | 0.94 | 0.47 | 0.77 | 0.11 | 0.88 | 1.49 | 0.74 |
| **Autobiographical interview** |  |  |  |  |  |  |  |  |
| Internal details | 0.06 | 0.98 | 0.48 | 0.92 | 0.21 | 0.97 | 1.51 | 0.92 |
| Internal events | 1.30 | 0.92 | 1.74 | 0.92 | 0.45 | 0.92 | 1.34 | 0.92 |
| Internal time | 3.76 | 0.92 | 3.44 | 0.92 | 2.43 | 0.92 | 1.05 | 0.92 |
| Internal place | 0.15 | 0.97 | 0.09 | 0.97 | 0.20 | 0.97 | 0.00 | 0.99 |
| Internal perceptual | 0.54 | 0.92 | 0.73 | 0.92 | 0.19 | 0.97 | 0.70 | 0.92 |
| Internal thoughts/emotions | 1.06 | 0.92 | 1.25 | 0.92 | 0.28 | 0.93 | 0.56 | 0.92 |
| Vividness ratings | 2.16 | 0.92 | 1.17 | 0.92 | 2.12 | 0.92 | 0.00 | 0.99 |
| **Future thinking** |  |  |  |  |  |  |  |  |
| Experiential index | 0.87 | 0.76 | 0.97 | 0.76 | 0.42 | 0.82 | 0.50 | 0.82 |
| Spatial references | 2.45 | 0.59 | 1.95 | 0.59 | 2.07 | 0.59 | 0.37 | 0.82 |
| Entities present | 0.38 | 0.82 | 0.31 | 0.84 | 0.27 | 0.84 | 0.03 | 0.95 |
| Sensory descriptions | 0.38 | 0.82 | 0.40 | 0.82 | 0.20 | 0.84 | 0.29 | 0.84 |
| Thoughts/emotions/actions | 0.07 | 0.90 | 0.15 | 0.88 | 0.00 | 0.98 | 0.22 | 0.84 |
| Spatial coherence index | 0.16 | 0.87 | 0.42 | 0.82 | 0.00 | 0.98 | 0.41 | 0.82 |
| **Navigation** |  |  |  |  |  |  |  |  |
| Overall navigation score | 0.87 | 0.73 | 1.48 | 0.68 | 0.07 | 0.91 | 1.82 | 0.62 |
| Scene recognition | 0.02 | 0.92 | 0.06 | 0.91 | 0.00 | 1.00 | 0.08 | 0.91 |
| Proximity judgements | 1.34 | 0.68 | 0.49 | 0.80 | 1.95 | 0.62 | 0.08 | 0.91 |
| Route knowledge | 0.72 | 0.76 | 1.72 | 0.63 | 0.00 | 0.99 | 2.63 | 0.50 |
| Sketch map | 0.20 | 0.89 | 0.00 | 0.99 | 0.87 | 0.73 | 0.60 | 0.76 |

P values are Benjamini-Hochberg false discovery rate corrected at $p < 0.05$ .

**Table S34.** Comparison of hippocampal grey matter <u>R_1_</u> when taking the <u>best and worst</u> performing participants for task, with age, gender, total intracranial volume and MRI scanner included as covariates.

| Performance variable | Whole hippocampus |  | Anterior hippocampus |  | Posterior hippocampus |  | Posterior/Anterior hippocampus ratio |  |
| --- | --- | --- | --- | --- | --- | --- | --- | --- |
|  | F | p | F | p | F | p | F | p |
| **Scene construction** |  |  |  |  |  |  |  |  |
| Experiential index | 1.43 | 0.74 | 2.25 | 0.74 | 0.74 | 0.74 | 1.60 | 0.74 |
| Spatial references | 2.89 | 0.62 | 3.62 | 0.62 | 1.91 | 0.74 | 0.23 | 0.82 |
| Entities present | 0.08 | 0.88 | 0.00 | 0.98 | 0.23 | 0.82 | 0.88 | 0.74 |
| Sensory descriptions | 0.76 | 0.74 | 1.53 | 0.74 | 0.33 | 0.79 | 0.64 | 0.74 |
| Thoughts/emotions/actions | 0.41 | 0.78 | 0.93 | 0.74 | 0.12 | 0.88 | 3.63 | 0.62 |
| Spatial coherence index | 1.23 | 0.74 | 1.09 | 0.74 | 1.21 | 0.74 | 0.00 | 0.98 |
| **Autobiographical interview** |  |  |  |  |  |  |  |  |
| Internal details | 2.41 | 0.92 | 1.66 | 0.92 | 2.40 | 0.92 | 0.12 | 0.97 |
| Internal events | 3.22 | 0.92 | 1.70 | 0.92 | 4.21 | 0.92 | 0.84 | 0.92 |
| Internal time | 0.05 | 0.99 | 0.33 | 0.93 | 0.00 | 0.99 | 0.43 | 0.92 |
| Internal place | 0.58 | 0.92 | 0.81 | 0.92 | 0.38 | 0.92 | 0.40 | 0.92 |
| Internal perceptual | 3.99 | 0.92 | 5.06 | 0.92 | 2.74 | 0.92 | 0.66 | 0.92 |
| Internal thoughts/emotions | 0.42 | 0.92 | 0.07 | 0.97 | 0.69 | 0.92 | 0.45 | 0.92 |
| Vividness ratings | 0.57 | 0.92 | 0.31 | 0.93 | 0.72 | 0.92 | 0.12 | 0.97 |
| **Future thinking** |  |  |  |  |  |  |  |  |
| Experiential index | 2.04 | 0.59 | 2.57 | 0.59 | 1.24 | 0.73 | 0.22 | 0.84 |
| Spatial references | 8.47 | 0.25 | 7.62 | 0.25 | 8.00 | 0.25 | 0.00 | 0.98 |
| Entities present | 6.57 | 0.30 | 6.04 | 0.30 | 5.08 | 0.30 | 0.38 | 0.82 |
| Sensory descriptions | 3.00 | 0.59 | 2.44 | 0.59 | 2.79 | 0.59 | 0.03 | 0.95 |
| Thoughts/emotions/actions | 1.09 | 0.73 | 0.23 | 0.84 | 1.82 | 0.59 | 2.23 | 0.59 |
| Spatial coherence index | 0.71 | 0.82 | 0.66 | 0.82 | 0.63 | 0.82 | 0.01 | 0.98 |
| **Navigation** |  |  |  |  |  |  |  |  |
| Overall navigation score | 0.03 | 0.92 | 0.04 | 0.92 | 0.18 | 0.89 | 0.91 | 0.73 |
| Scene recognition | 0.06 | 0.91 | 0.25 | 0.88 | 0.00 | 0.99 | 0.65 | 0.76 |
| Proximity judgements | 1.35 | 0.68 | 0.89 | 0.73 | 1.31 | 0.68 | 0.07 | 0.91 |
| Route knowledge | 0.43 | 0.80 | 1.06 | 0.73 | 0.10 | 0.91 | 1.34 | 0.68 |
| Sketch map | 3.97 | 0.48 | 3.54 | 0.48 | 3.62 | 0.48 | 0.06 | 0.91 |

P values are Benjamini-Hochberg false discovery rate corrected at $p < 0.05$ .

**Table S35.** Comparison of hippocampal grey matter <u>R_2_*</u> when taking the <u>best and worst</u> performing participants for task, with age, gender, total intracranial volume and MRI scanner included as covariates.

| Performance variable | Whole hippocampus |  | Anterior hippocampus |  | Posterior hippocampus |  | Posterior/Anterior hippocampus ratio |  |
| --- | --- | --- | --- | --- | --- | --- | --- | --- |
|  | F | p | F | p | F | p | F | p |
| **Scene construction** |  |  |  |  |  |  |  |  |
| Experiential index | 5.19 | 0.56 | 4.04 | 0.62 | 4.24 | 0.62 | 0.09 | 0.88 |
| Spatial references | 3.02 | 0.62 | 0.46 | 0.77 | 5.61 | 0.56 | 1.98 | 0.74 |
| Entities present | 1.03 | 0.74 | 0.23 | 0.82 | 1.78 | 0.74 | 0.61 | 0.74 |
| Sensory descriptions | 0.51 | 0.77 | 0.00 | 0.98 | 1.55 | 0.74 | 1.87 | 0.74 |
| Thoughts/emotions/actions | 0.02 | 0.96 | 0.17 | 0.85 | 0.27 | 0.82 | 0.89 | 0.74 |
| Spatial coherence index | 0.01 | 0.98 | 0.67 | 0.74 | 1.02 | 0.74 | 3.23 | 0.62 |
| **Autobiographical interview** |  |  |  |  |  |  |  |  |
| Internal details | 0.40 | 0.92 | 0.12 | 0.97 | 0.61 | 0.92 | 0.15 | 0.97 |
| Internal events | 0.45 | 0.92 | 0.08 | 0.97 | 0.86 | 0.92 | 0.63 | 0.92 |
| Internal time | 2.30 | 0.92 | 2.27 | 0.92 | 1.08 | 0.92 | 0.29 | 0.93 |
| Internal place | 0.30 | 0.93 | 0.75 | 0.92 | 0.07 | 0.97 | 0.60 | 0.92 |
| Internal perceptual | 2.91 | 0.92 | 4.19 | 0.92 | 0.96 | 0.92 | 2.09 | 0.92 |
| Internal thoughts/emotions | 0.03 | 0.99 | 0.07 | 0.97 | 0.37 | 0.92 | 0.75 | 0.92 |
| Vividness ratings | 0.95 | 0.92 | 0.37 | 0.92 | 1.01 | 0.92 | 0.08 | 0.97 |
| **Future thinking** |  |  |  |  |  |  |  |  |
| Experiential index | 5.16 | 0.30 | 2.96 | 0.59 | 5.15 | 0.30 | 0.20 | 0.84 |
| Spatial references | 2.02 | 0.59 | 2.16 | 0.59 | 1.24 | 0.73 | 0.13 | 0.88 |
| Entities present | 4.31 | 0.39 | 2.53 | 0.59 | 5.71 | 0.30 | 0.09 | 0.90 |
| Sensory descriptions | 1.91 | 0.59 | 1.42 | 0.71 | 1.76 | 0.60 | 0.05 | 0.94 |
| Thoughts/emotions/actions | 1.82 | 0.59 | 0.90 | 0.76 | 2.43 | 0.59 | 0.53 | 0.82 |
| Spatial coherence index | 0.47 | 0.82 | 1.38 | 0.71 | 0.00 | 1.00 | 1.16 | 0.73 |
| **Navigation** |  |  |  |  |  |  |  |  |
| Overall navigation score | 0.24 | 0.88 | 0.20 | 0.89 | 1.98 | 0.62 | 4.93 | 0.48 |
| Scene recognition | 2.19 | 0.58 | 0.59 | 0.76 | 3.07 | 0.50 | 0.60 | 0.76 |
| Proximity judgements | 0.03 | 0.92 | 0.11 | 0.91 | 0.81 | 0.75 | 0.31 | 0.86 |
| Route knowledge | 0.42 | 0.80 | 0.16 | 0.90 | 2.97 | 0.50 | 3.75 | 0.48 |
| Sketch map | 0.42 | 0.80 | 2.23 | 0.58 | 0.25 | 0.88 | 4.54 | 0.48 |

P values are Benjamini-Hochberg false discovery rate corrected at $p < 0.05$ .

